# Differential Regulation of Retinoic Acid Metabolism in Fanconi Anemia

**DOI:** 10.1101/2023.04.06.535759

**Authors:** Justin L. Blaize, Bahaa M. Noori, Kelsey P. Hunter, Kathryn A. Henrikson, Janet A. Atoyan, Alan A. Ardito, Frank X. Donovan, Settara C. Chandrasekharappa, Detlev Schindler, Niall G. Howlett

**Author notes:** Corresponding author: Niall G. Howlett Ph.D., 379 Center for Biotechnology and Life Sciences, 120 Flagg Road, Kingston, RI, USA.

## Abstract

Fanconi anemia (FA) is a rare genetic disease characterized by heterogeneous congenital abnormalities and increased risk for bone marrow failure and cancer. FA is caused by mutation of any one of 23 genes, the protein products of which function primarily in the maintenance of genome stability. An important role for the FA proteins in the repair of DNA interstrand crosslinks (ICLs) has been established *in vitro*. While the endogenous sources of ICLs relevant to the pathophysiology of FA have yet to be clearly determined, a role for the FA proteins in a two-tier system for the detoxification of reactive metabolic aldehydes has been established. To discover new metabolic pathways linked to FA, we performed RNA-seq analysis on non-transformed FA-D2 (*FANCD2^-/-^*) and FANCD2-complemented patient cells. Multiple genes associated with retinoic acid metabolism and signaling were differentially expressed in FA-D2 (*FANCD2^-/-^*) patient cells, including *ALDH1A1* and *RDH10*, which encode for retinaldehyde and retinol dehydrogenases, respectively. Increased levels of the ALDH1A1 and RDH10 proteins was confirmed by immunoblotting. FA-D2 (*FANCD2^-/-^*) patient cells displayed increased aldehyde dehydrogenase activity compared to the FANCD2-complemented cells. Upon exposure to retinaldehyde, FA-D2 (*FANCD2^-/-^*) cells exhibited increased DNA double-strand breaks and checkpoint activation indicative of a defect in the repair of retinaldehyde-induced DNA damage. Our findings describe a novel link between retinoic acid metabolism and FA and identify retinaldehyde as an additional reactive metabolic aldehyde relevant to the pathophysiology of FA.

## INTRODUCTION

Fanconi anemia (FA) is a rare genetic disease, characterized by congenital abnormalities, increased risk for bone marrow failure and both hematologic and nonhematologic cancers, accelerated aging, and premature mortality (1). Most FA patients harbor compound heterozygous germline mutations in one of twenty-three genes. Consequently, age at diagnosis and clinical manifestations can vary widely. Long-term therapeutic options for FA patients remain underdeveloped. A greater understanding of the molecular etiology of FA is critical for the discovery of new therapeutic options for FA patients.

At the cellular level, the proteins encoded by the FA genes function collectively in the maintenance of genome stability. Exemplifying this function, FA patient cells frequently exhibit spontaneous structural and numerical chromosome aberrations (2,3). *In vitro* studies have established a critical role for the FA proteins in the repair of DNA interstrand crosslinks (ICLs), a complex process that encompasses elements of the canonical DNA repair pathways nucleotide excision repair (NER), translesion DNA synthesis (TLS), and homologous recombination (HR) (4). However, the endogenous sources of chromosome instability, including ICLs, in the physiological setting remain to be determined. Several studies over the past decade indicate that aldehydes generated during metabolic processes pose an acute genotoxic threat in the setting of FA. For example, mice with combined inactivation of *Fancd2* and *Aldh2* (which oxidizes acetaldehyde to acetate) exhibit developmental defects and increased risk for leukemia and/or aplastic anemia (5,6). Similarly, combined inactivation of the FA pathway and *Adh5* (major formaldehyde detoxifying enzyme) leads to synthetic lethality (7,8). These findings have led to the hypothesis that the FA pathway functions in a two-tier protection system to mitigate the cytotoxic and genotoxic effects of endogenous metabolic aldehydes. In the first tier, substrate-specific aldehyde dehydrogenases oxidize reactive aldehydes to their corresponding nontoxic carboxylic acids. In the second tier, aldehydes that escape oxidation and react with DNA are repaired in an FA-pathway dependent manner. Inactivation of both tiers leads to severe cellular and physiological consequences (9,10). It remains to be determined if this system extends to other metabolic aldehydes, in addition to acetaldehyde and formaldehyde.

Retinoid metabolism is the cellular process whereby vitamin A (retinol) from dietary sources is converted to all-*trans* retinoic acid (*at*RA), a signaling molecule critical for patterning, morphogenesis, and organogenesis during development (11,12). During *at*RA generation, retinol must first be oxidized to retinaldehyde, a reaction carried out by the ALDH1A1, ALDH1A2, and ALDH1A3 paralogous aldehyde dehydrogenases. As retinaldehyde is a potential genotoxin, vitamin A levels must be precisely regulated during pregnancy, as excessive or insufficient levels can lead to congenital abnormalities and/or embryonic lethality (13). Nuclear *at*RA functions primarily as a regulator of gene expression. *at*RA binds to heterodimers of two classes of nuclear receptors, the retinoic acid receptors (RARs) and the retinoid X receptors (RXRs). RAR-RXR bind to retinoic acid response elements (RAREs) in gene promoters and, upon *at*RA binding, recruit transcriptional co-activators and histone acetyltransferases (HATs) to activate gene expression. In the absence of *at*RA, the RAR-RXR nuclear receptors promote gene silencing *via* corepressor/histone deacetylase (HDAC) complex recruitment (14).

In this study, we describe a novel link between FA and retinoic acid metabolism. RNA- seq analysis of non-transformed FA-D2 (*FANCD2^-/-^*) and FANCD2-complemented patient cells revealed differential expression of multiple genes linked to retinoic acid metabolism and signaling in FA-D2 (*FANCD2^-/-^*) cells compared to FANCD2 complemented FA-D2 cells. For example, we observed markedly increased expression of the *ALDH1A1* and *RDH10* genes, which encode for retinaldehyde and retinol dehydrogenases, respectively, in FA-D2 (*FANCD2^-/-^*) cells. FA-D2 (*FANCD2^-/-^*) cells displayed increased aldehyde dehydrogenase activity compared to the FANCD2 complemented FA-D2 cells. Upon exposure to retinaldehyde, FA-D2 (*FANCD2^-/-^*) cells exhibited increased DNA double-strand breaks, replication arrest, and checkpoint activation indicative of a defect in the repair of retinaldehyde-induced DNA damage. Our findings describe a novel link between retinoic acid metabolism and FA and identify retinaldehyde as an additional reactive metabolic aldehyde linked to the pathophysiology of FA.

## RESULTS

### Characterization of a novel FA-D2 patient model

The FANCD2 protein has been well established to play a key role in DNA interstrand crosslink (ICL) repair and in the cellular DNA replication stress response (4,15). To discover additional cellular pathways linked to FA, and FANCD2, we sought to perform global transcriptional profiling of a non-transformed FA-D2 cell model. FA-D2 (*FANCD2^-/-^*) and FANCD2-complemented skin fibroblasts isolated from a 1-year-old patient with a severe clinical phenotype (Supplementary Table 1) were immortalized with human telomerase and characterized for functional complementation. Compound heterozygous mutations in the *FANCD2* gene were confirmed by Sanger sequencing; a G>A transition at nucleotide 2444 in exon 26 [c.2444G>A (exon 26)] leading to the missense change R815Q and a G>A transition at the first nucleotide of intron 28 [c.2715+1G>A (IVS28+1G>A)], leading to a frameshift and premature truncation E906Lfs*4 (Figure 1A). No FANCD2 expression was observed in the FA-D2 (*FANCD2^-/-^*) cells, while robust expression was observed in the FANCD2-complemented cells (Figure 1B). Compared to FANCD2-complemented cells, FA-D2 (*FANCD2^-/-^*) cells displayed increased chromosome breaks following exposure to the DNA interstrand crosslinking agents mitomycin C (MMC) and diepoxybutane (DEB) (D. Schindler and A. Smogorzewska, personal communication). Consistent with previous studies with non-transformed cells, and in contrast to that observed in transformed lines (16,17), we observed low levels of FANCD2 monoubiquitination following MMC treatment and treatment with the DNA polymerase inhibitor aphidicolin (APH) (Figure 1B). FA-D2 (*FANCD2^-/-^*) cells exhibited increased levels of CHK1 pS345, MRE11, and RPA2 following MMC and APH treatment compared to FANCD2-complemented cells, consistent with increased levels of DNA damage in these cells (Figure 1B). In a G2/M accumulation assay, FA-D2 (*FANCD2^-/-^*) cells demonstrated a 2-fold increase in G2/M phase cells compared to FANCD2-complemented cells following MMC treatment, consistent with a block in cell cycle progression in the absence of FANCD2 (Figure 1C). These results confirm the functional complementation of this non-transformed FA-D2 patient line.

**Figure 1.**
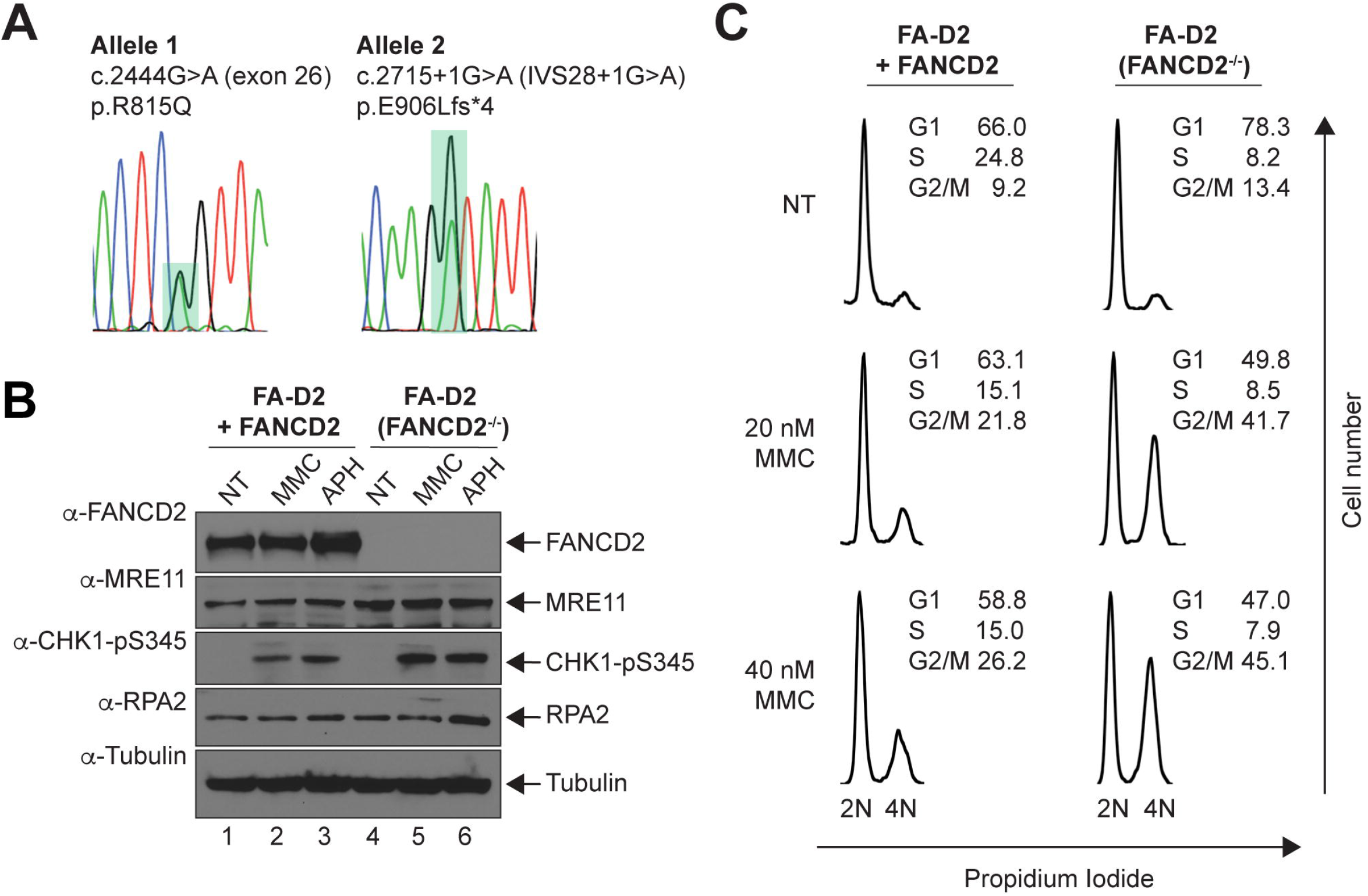
Characterization of a novel FA-D2 patient model. (A) Sanger sequencing confirmation of compound heterozygous mutations in the *FANCD2* gene in ACHT (FA-D2) patient fibroblasts; a G>A transition at the first nucleotide of intron 28 [c.2715+1G>A (IVS28+1G>A)], leading to a frameshift and premature truncation E906Lfs*4 and a G>A transition at nucleotide 2444 in exon 26 [c.2444G>A (exon 26)] leading to the missense change R815Q. (B) FA-D2 (*FANCD2^-/-^*) and FANCD2-complemented FA-D2 cells were incubated in the absence and presence of 200 nM mitomycin C (MMC) or 1 µM aphidicolin (APH) for 24 h and whole cell lysates were analyzed by immunoblotted using anti-FANCD2, anti-MRE11, anti-RPA2, and anti-CHK1 pS345 antibodies. (C) FA-D2 (*FANCD2^-/-^*) and FANCD2-complemented FA-D2 cells were incubated in the absence or presence of MMC (20 nM and 40 nM) MMC for 24 hours, cells were fixed and stained with propidium iodide, and analyzed by flow cytometry.

### RNA-seq analysis reveals several differentially expressed gene sets in FA-D2 (*FANCD2^-/-^*) cells

RNA-seq analysis was performed with FA-D2 (*FANCD2^-/-^*) and FANCD2-complemented FA-D2 cells. Quadruplicate biological replicates were sequenced at a read depth of 60M reads per sample. Principal component analysis (PCA) revealed clustering by genotype with 85% of the variance attributed to gene expression differences between the FA-D2 (*FANCD2^-/-^*) and the FANCD2-complemented cells (Figure 2A). A clustered heatmap illustrates the differential gene expression profiles between mutant and complemented FA-D2 (*FANCD2^-/-^*) cells (Figure 2B). Our analysis uncovered 1,458 significantly differentially expressed genes (adjusted P value (Padj) < 0.05 and a log2 fold change greater than 1) between FA-D2 (*FANCD2^-/-^*) and FANCD2- complemented cells (Figure 2C). Significantly differentially expressed genes were clustered by gene ontology and the enrichment of gene ontology terms was tested using Fisher’s exact test. Gene ontology terms significantly enriched with a Padj value < 0.05 in the differentially expressed gene sets included extracellular matrix organization (GO:0030198), positive regulation of gene expression (GO:0010628), angiogenesis (GO:0001525), nervous system development (GO:0007399), and the cellular response to retinoic acid (GO:0071300). In addition, gene set enrichment analysis (GSEA) demonstrates enrichment for genes with defined roles in angiogenesis, neurogenesis, and the cellular response to retinoic acid (Figure 2D). The cellular response to retinoic acid/retinoic acid metabolism was of particular interest given recent established connections between the FA pathway and the detoxification of reactive metabolic aldehydes (9,10).

**Figure 2.**
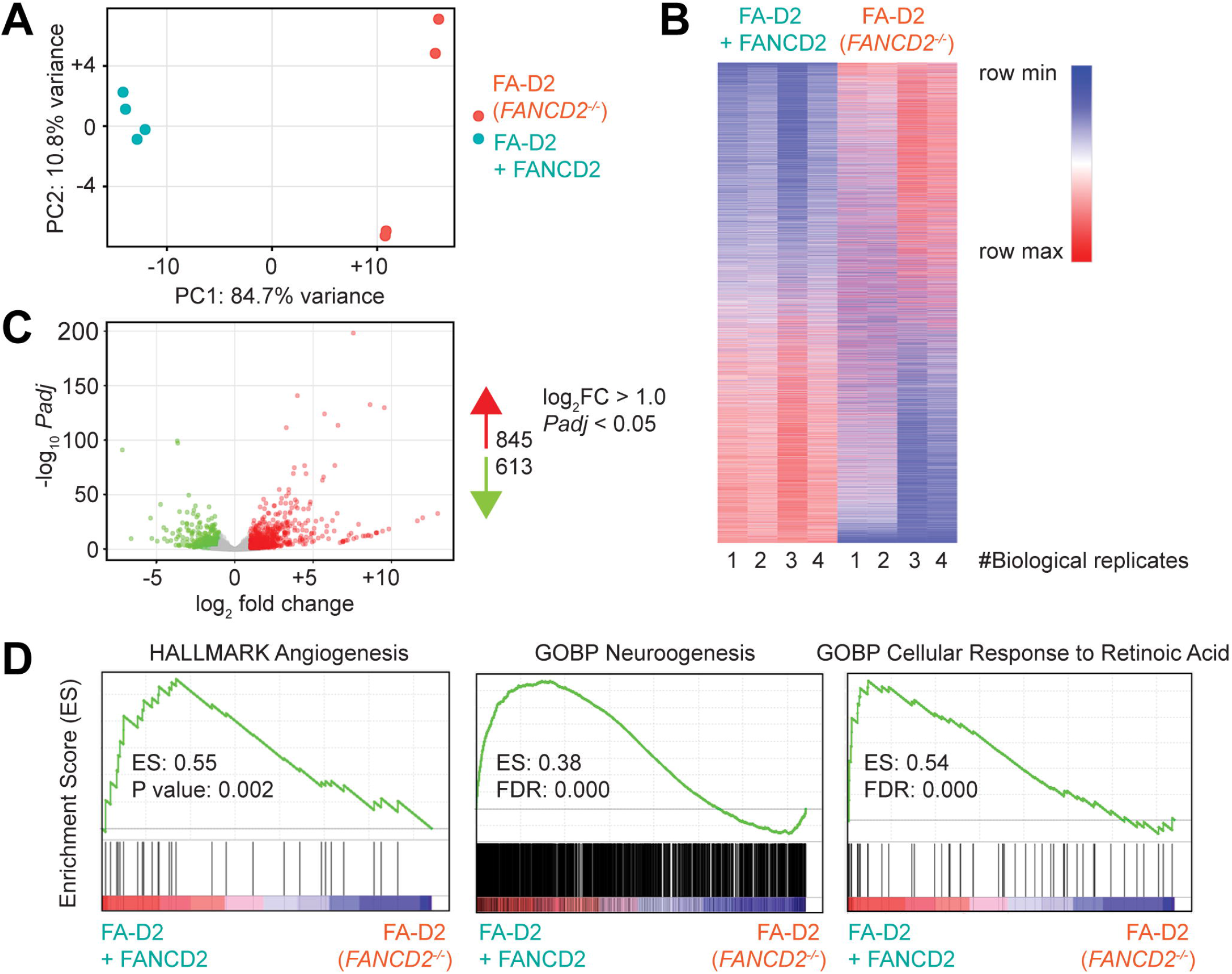
RNA-seq analysis reveals several differentially expressed gene between FA-D2 (*FANCD2^-/-^*) and FANCD2-complemented cells. (A) Principal component analysis (PCA) of quadruplicate biological samples were projected onto a 2D plane spanned by their first two principal components. The percentage of the total variance per direction is shown. The x-axis depicts the most variance and the y-axis depicts the second most. (B) RNA-seq heatmap of differentially expressed genes between FA-D2 (*FANCD2^-/-^*) and FANCD2-complemented FA- D2 cells. Up-regulated genes are shown in red and down-regulated genes are shown in blue. (C) Volcano plot of differentially expressed genes between FA-D2 (*FANCD2^-/-^*) and FANCD2- complemented FA-D2 cells. Genes with an adjusted p-value less than 0.05 and a log2 fold change greater than 1 are shown in red while genes with an adjusted p-value less than 0.05 and a log2 fold change less than -1 are shown in green. (D) RNA-seq expression data of FA-D2 (*FANCD2^-/-^*) and FANCD2-complemented FA-D2 cells were further analyzed using gene set enrichment analysis (GSEA) (v4.2.1) platform sourced from Broad Institute, for angiogenesis, neurogenesis, and cellular response to retinoic acid gene sets.

### Dysregulation of retinoic acid metabolism in non-transformed FA-D2 (*FANCD2^-/-^*) cells

RNA-seq analysis uncovered differential expression of multiple genes linked to retinoic acid metabolism and/or retinoic acid signaling in FA-D2 (*FANCD2^-/-^*) compared to FANCD2- complemented FA-D2 cells (Figure 3A and Supplementary Table 2). For example, we observed a negative 7.2 log2 fold change of the *ALDH1A1* gene, indicating markedly increased expression in FA-D2 (*FANCD2^-/-^*) cells compared to FANCD2-complemented cells. *ALDH1A1* was the most highly differentially expressed *ALDH* gene, out of 15 of the 19 human ALDH genes expressed in these cells (Supplementary Table 3). *ALDH1A1* encodes for an aldehyde dehydrogenase that irreversibly oxidizes retinaldehyde to retinoic acid. We also observed negative 1.4 and 1.1 log2 fold changes for *RDH10* and *CRABP2*, respectively (Figure 3A and Supplementary Table 2). *RDH10* encodes for retinol dehydrogenase and converts retinol to retinaldehyde in a reversible reaction. *CRABP2* encodes for cellular retinoic acid binding protein II and transports cytoplasmic retinoic acid to the nucleus (11,12). A very minor difference in *ALDH2* gene expression was observed (-0.6 log2 fold change), while we detected a -1.1 log2 fold change for the *ADH5* gene (Supplementary Tables 3 and 4). We used immunoblotting to confirm the increased expression of ALDH1A1, RDH10, and CRABP2 in the FA-D2 (*FANCD2^-/-^*) cells using positive and negative controls (Figure 3B). A549 cells, a lung epithelial-like carcinoma line, were used as a positive control for ALDH1A1 expression as this line is known to express high levels of this protein (Human Protein Atlas) (18,19). Increased ALDH1A1 expression in FA-D2 (*FANCD2^-/-^*) cells was also confirmed using the isoform-selective fluorescent ALDH1A1 probe, AlDeSense (Figure 3C) (20). Cellular fractionation experiments revealed that both ALDH1A1 and RDH10 expression were confined to the cytoplasm, with no change in subcellular localization observed following expsoure of the cells to exogenous retinaldehyde (Supplementary Figure S1A). While CRABP2 expression was primarily cytoplasmic, we did observe low levels of expression in the nucleus consistent with its retinoic acid shuttling function (Supplementary Figure S1A). We next analyzed ALDH1A1 and ALDH2 expression in FA-A (*FANCA^-/-^*), FA-F (*FANCF^-/-^*), and FA-D2 (*FANCD2^-/-^*) patient lymphoblasts and their FANCA-, FANCF-, or FANCD2-complemented counterparts, respectively (Supplementary Figure S1B). In contrast to the non-transformed FA-D2 (*FANCD2^-/-^*) fibroblasts, we did not observe ALDH1A1 expression in any of the lymphoblasts tested (Supplementary Figure S1B). We did observe increased expression of ALDH2, however, in FA-A, FA-F, and FA-D2 lymphoblasts, compared to their complemented counterparts (Supplementary Figure S1B). We also examined Aldh1a1 and Rdh10 expression in wild-type, *Fancd2^+/+^/Aldh2^-/-^*, *Fancd2^-/-^/Aldh2^+/+^,* and *Fancd2^-/-^/Aldh2^-/-^*non-transformed mouse ear fibroblasts. In contrast to wild-type cells, we observed an Aldh1a1 immunoreactive band - albeit at a lower molecular weight than expected - and increased levels of Rdh10 in *Fancd2^+/+^/Aldh2^-/-^*, *Fancd2^-/-^/Aldh2^+/+^,* and *Fancd2^-/-^/Aldh2^-/-^*cells (Supplementary Figure S1C). We also examined Aldh1a1 and Rdh10 protein expression in *Fancd2^+/+^*, *Fancd2^+/-^,* and *Fancd2^-/-^* mouse embryonic fibroblasts (MEFs) and observed increased levels of Aldh1a1 in *Fancd2^-/-^* MEFs compared to *Fancd2^+/+^*and *Fancd2^+/-^* MEFs (Supplementary Figure S1D). No appreciable differences in Rdh10 expression levels between these cells were observed (Supplementary Figure S1D). To determine if increased ALDH1A1 expression results in higher aldehyde dehydrogenase (ALDH) activity in the non-transformed FA-D2 (*FANCD2^-/-^*) cells, we performed an ALDEFLUOR assay with FA-D2 (*FANCD2^-/-^*) and FANCD2-complemented cells. Cells were incubated with the ALDH substrate BODIPY-aminoacetaldehyde (BAAA), which is converted into the fluorescent molecule BODIPY-aminoacetate (BAA) *via* ALDH activity. This reaction was performed in the absence and presence of the ALDH inhibitor N,N-diethylaminobenzaldehyde (DEAB). FA-D2 (*FANCD2^-/-^*) cells showed an approximate 2-fold increase in ALDH activity compared to FANCD2-complemented cells, indicating that increased ALDH1A1 expression is associated with higher ALDH activity in these cells (Figure 3C). HeLa and A549 cells were included as negative and positive controls for ALDH activity, respectively (Figure 3C).

**Figure 3.**
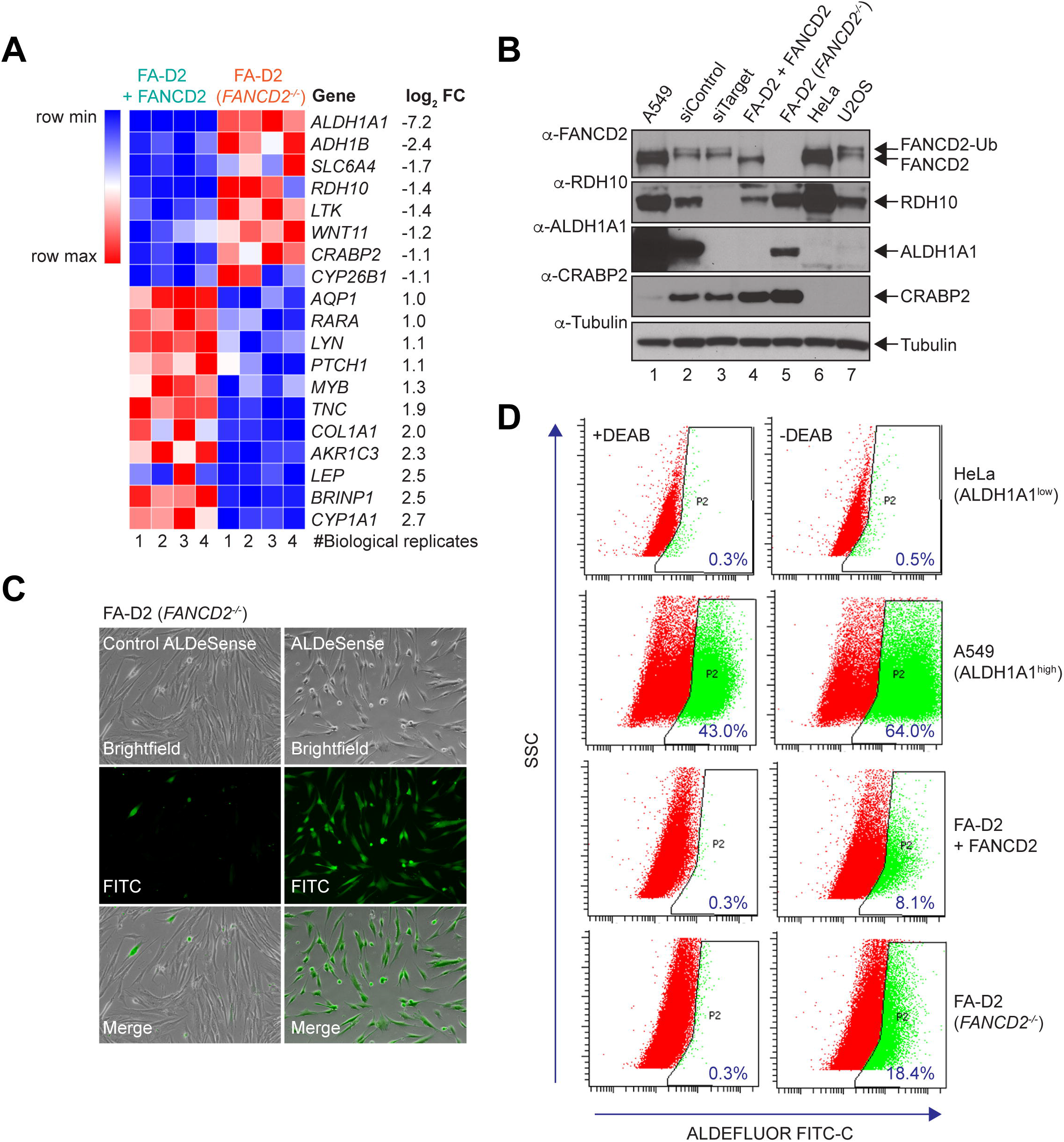
Differential regulation of retinoic acid metabolism and signaling in FA-D2 (*FANCD2^-/-^*) patient cells. (A) Morpheus heatmap analysis demonstrating differential expression of retinoic acid metabolism and signaling genes in FA-D2 (*FANCD2^-/-^*) cells relative to FANCD2-complemented FA-D2 cells. Genes with increased expression in FA-D2 (*FANCD2^-/-^*) cells relative to FANCD2-complemented FA-D2 cells are depicted in red and genes with decreased expression are depicted in blue. (B) Immunoblotting validation of increased expression of ALDH1A1 and RDH10 in FA-D2 (*FANCD2^-/-^*) cells relative to FANCD2- complemented FA-D2 cells. Antibody-protein specificity was confirmed by depleting cells of ALDH1A1 or RDH10 using pooled siRNAs (lane 3). Whole-cell lysates were prepared from the indicated cells and immunoblotted using the indicated antibodies. (C) ALDEFLUOR flow cytometry assay demonstrating increased aldehyde dehydrogenase (ALDH) activity in FA-D2 (*FANCD2^-/-^*) cells compared to FANCD2-complemented FA-D2 cells. ALDH-negative HeLa cells and ALDH-positive A549 cells were used as negative and positive controls, respectively. Cells were incubated in the absence or presence of the ALDH inhibitor N,N-diethylaminobenzaldehyde (DEAB) prior to analysis.

### Increased retinaldehyde genotoxicity in FA patient cells

Aldehydes are strong electrophiles, and the terminal carbonyl group can form adducts with various cellular targets including DNA. The mechanisms of retinaldehyde genotoxicity have not been clearly elucidated. To learn more about the mechanisms of retinaldehyde genotoxicity, we exposed HeLa and U2OS cells to the type II topoisomerase inhibitor etoposide (VP-16) and all-*trans*-retinaldehyde (a*t*RALD) and analyzed protein biomarkers of DNA damage, DNA repair, and cell cycle checkpoint activation.

As expected, VP-16 exposure led to a marked increase in H2AX S139 (*γ*H2AX) and CHK1 S345 phosphorylation, indicative of DNA double-strand break formation and activation of the intra-S phase checkpoint, respectively (Figures 4A and B). At later time points following exposure, atRALD treatment also resulted in increased levels of *γ*H2AX, albeit to a lesser extent than VP- 16 (Figures 4A and B). While a*t*RALD exposure led to a modest increase in CHK1 S345 phosphorylation in U2OS cells, induction of CHK2 T68 phosphorylation was observed in both HeLa and U2OS, suggesting that a*t*RALD exposure activates the G2-M phase checkpoint (Figures 4A and B). As ALDH1A1 exhibits specificity for a*t*RALD and 9-*cis*-retinaldehyde (9*c*RALD), we also exposed HeLa and U2OS cells to both isomers and analyzed the same protein biomarkers of DNA damage, DNA repair, and cell cycle checkpoint activation. Exposure to both a*t*RALD and 9*c*RALD resulted in DNA damage and checkpoint activation to varying degrees. Under the conditions tested, however, we observed a greater increase in levels of *γ*H2AX following *at*RALD treatment, compared to 9*c*RALD (Supplementary Figure S2A and B). We next compared the response of the non-transformed FA-D2 (*FANCD2^-/-^*) and FANCD2- complemented cells to a*t*RALD. We observed increased levels of *γ*H2AX and CHK2 T68 phosphorylation in a*t*RALD-treated FA-D2 (*FANCD2^-/-^*) cells compared to the FANCD2- complemented cells (Figure 4C). Consistent with a*t*RALD activating the G2-M phase checkpoint, we observed reduced numbers of G2-M phase FA-D2 (*FANCD2^-/-^*) cells both with and without a*t*RALD exposure (Supplementary Figure S2C). We also observed increased *γ*H2AX nuclear foci formation in FA-D2 (*FANCD2^-/-^*) cells compared to the FANCD2- complemented cells following exposure to either a*t*RALD or VP-16 (Figure 4D and Supplementary Figure S2D). These results indicate that FA-D2 (*FANCD2^-/-^*) cells are compromised in their ability to repair a*t*RALD-induced DNA damage. Similar findings were observed for EBV-immortalized FA-A (*FANCA^-/-^*) cells, whereby a*t*RALD exposure led to increased *γ*H2AX and CHK2 T68 phosphorylation in FA-A (*FANCA^-/-^*) cells compared to FANCA-complemented cells (Supplementary Figure S2E and F). Taken together, these results suggest an important role for the FA pathway in the repair of *at*RALD-induced DNA damage, similar to that previously observed for acetaldehyde and formaldehyde.

**Figure 4.**
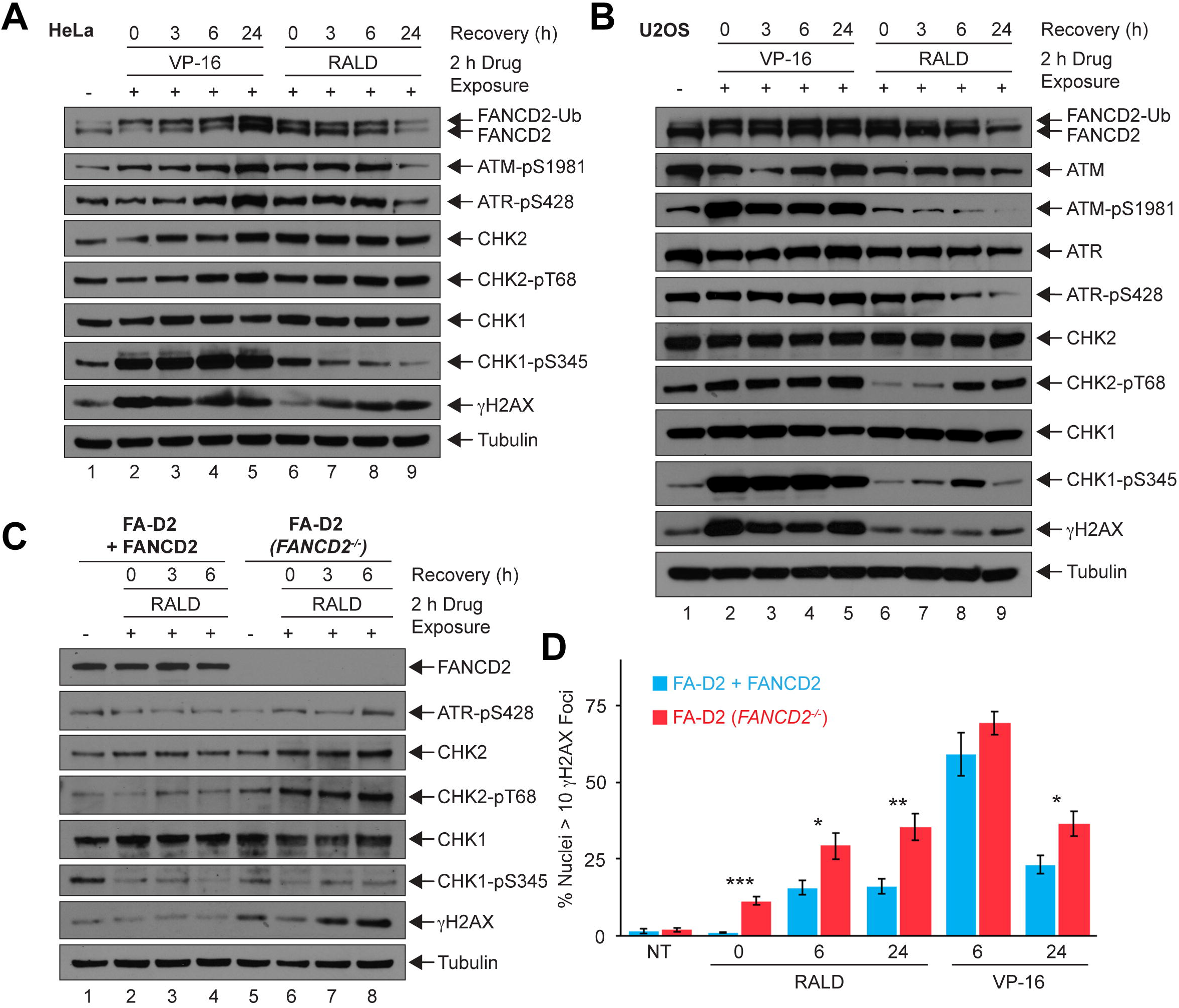
Increased retinaldehyde genotoxicity in FA-D2 (*FANCD2^-/-^*) patient cells. HeLa (A) and U2OS (B) cells were exposed to either 25 μM all-*trans*-retinaldehyde (a*t*RALD) or 25 μM etoposide (VP-16) for 2 h, released into drug-free media, and whole-cell lysates were prepared at 0, 3, 6, and 24 h following the exposure period. Whole-cell lysates were immunoblotted with the indicated antibodies. (C) Non-transformed FA-D2 (*FANCD2^-/-^*) and FANCD2-complemented cells were exposed to 25 μM a*t*RALD for 2 h, released into a*t*RALD- free media, and whole-cell lysates were prepared at 0, 3, and 6 h following the exposure period. Whole-cell lysates were immunoblotted with the indicated antibodies. (D) Non-transformed FA- D2 (*FANCD2^-/-^*) and FANCD2-complemented cells were exposed to 25 μM a*t*RALD or 25 μM VP-16 for 2 h and released into drug-free media. Immunofluorescence microscopy for *γ*H2AX was performed at the indicated time points following release. Error bars represent the standard errors of the means from two independent experiments. At least 600 nuclei were scored per biological replicate. ***, *P* < 0.001; **, *P* < 0.01; *, *P* < 0.05.

### Attenuated apoptosis in non-transformed FA-D2 (*FANCD2^-/-^*) cells

Gene set enrichment analysis (GSEA) of our RNA-seq data also revealed differential expression of apoptosis genes, suggesting downregulation of apoptosis in FA-D2 (*FANCD2^-/-^*) cells compared to the FANCD2- complemented cells (Figure 5A). To experimentally validate differential apoptosis gene expression, we analyzed cleavage of the poly(ADP)ribose polymerase enzyme PARP1 by immunofluorescence microscopy (IF). We observed an approximate 2-fold reduction in PARP1- positive nuclei following staurosporine (STS) exposure in FA-D2 (*FANCD2^-/-^*) cells compared to cells complemented with wild-type FANCD2 (Figure 5B). Similarly, using immunoblotting, we observed reduced STS-induced PARP, caspase 3 (CASP3), and caspase 7 (CASP7) cleavage in FA-D2 (*FANCD2^-/-^*) cells compared to FANCD2-complemented cells (Figure 5B). These results demonstrate that apoptosis is functionally attenuated in the non-transformed FA-D2 (*FANCD2^-/-^*) patient cells.

**Figure 5.**
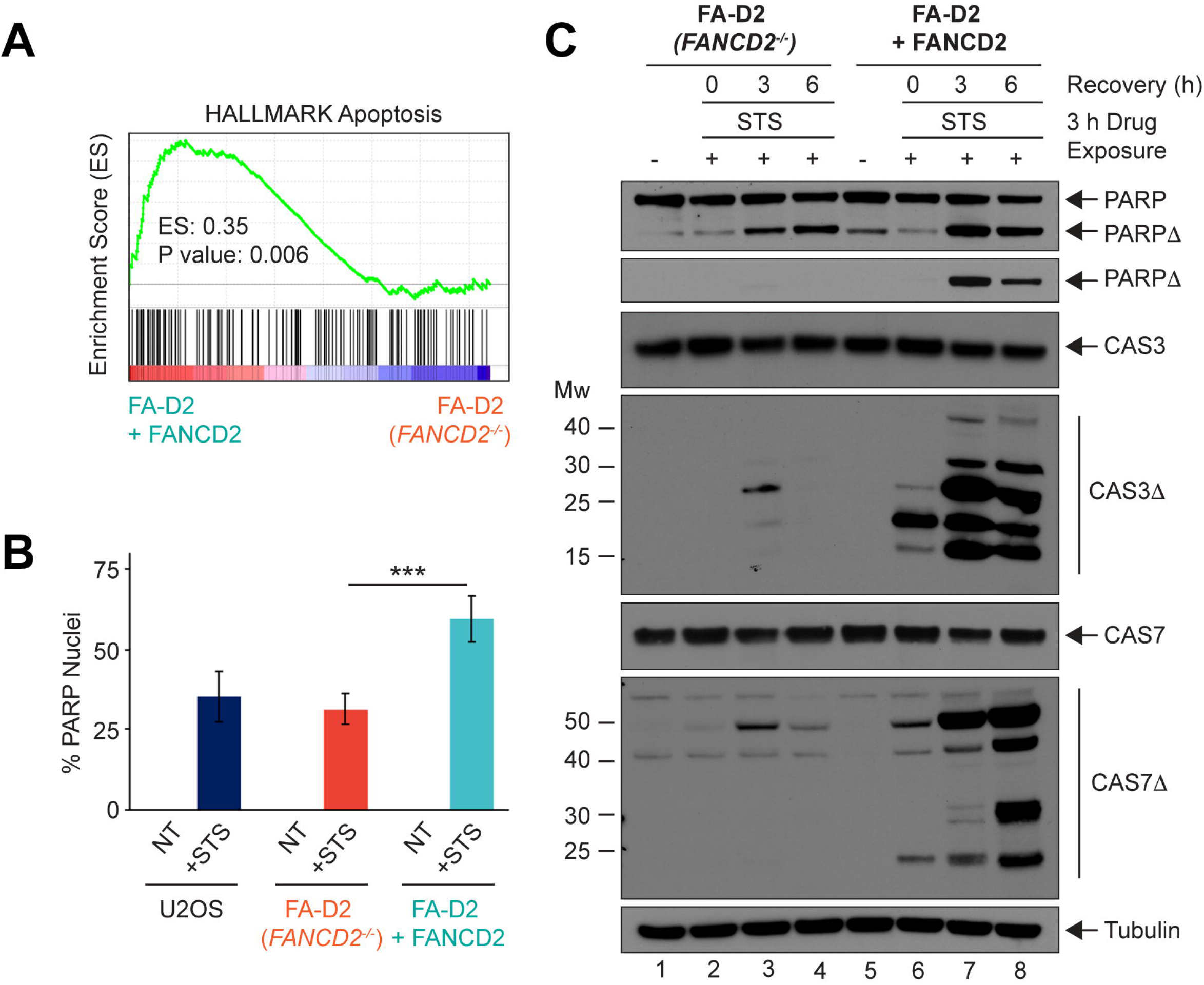
Attenuated apoptosis in non-transformed FA-D2 (*FANCD2^-/-^*) patient cells. (A) RNA-seq gene set enrichment analysis demonstrates increased expression of apoptosis genes in FANCD2-complemented cells compared to FA-D2 (*FANCD2^-/-^*) cells. (B) Increased PARP1 nuclear staining in FANCD2-complemented cells compared to FA-D2 (*FANCD2^-/-^*) cells. Cells were incubated in the absence or presence of 1 μM staurosporine (STS) for 3 h, released into STS-free media, and the numbers of PARP1-stained nuclei scored. Error bars represent the standard deviations from at least 5 technical replicate measurements from a single experiment. At least 500 nuclei were scored per sample. This experiment was repeated three times with similar findings. ***, *P* < 0.001. (C) Decreased STS-induced apoptosis in FA-D2 (*FANCD2^-/-^*) cells. FA-D2 (*FANCD2^-/-^*) and FANCD2-complemented cells were incubated in the absence or presence of 1 μM STS for 3 h, released into STS-free media, whole-cell lysates prepared at 0, 3, and 6 h, and immunoblotted with the indicated antibodies.

## DISCUSSION

In this manuscript, we describe a previously undiscovered molecular link between FA and retinoic acid metabolism and signaling. Bulk RNA-seq analysis of non-transformed FA-D2 patient cells and the same cells complemented with wild-type FANCD2 uncovered differential expression of multiple retinoic acid metabolism and signaling genes. For example, we observed markedly increased cytoplasmic expression of the *ALDH1A1* and *RDH10* genes in FA-D2 (*FANCD2^-/-^*) patient cells compared to FANCD2-complemented cells, findings confirmed by qPCR and immunoblotting. Consistent with increased expression of ALDH1A1, FA-D2 (*FANCD2^-/-^*) patient cells demonstrated increased aldehyde dehydrogenase activity as determined by the ALDEFLUOR™ assay. We did not detect ALDH1A1 expression in EBV-immortalized FA-A, FA-F, or FA-D2 patient lymphoblasts or SV40-transformed FA-A and FA-D2 fibroblasts. To the best of our knowledge, all these cells were derived from FA patients greater than 7 years of age. We did however observe increased levels of Aldh1a1 and Rdh10 in non-transformed mouse ear fibroblasts in the absence of Fancd2 or Aldh2 and increased Aldh1a1 in *Fancd2^-/-^* mouse embryonic fibroblasts. These results suggest that dysregulated retinoic acid metabolism might be a feature of non-transformed FA models and restricted to younger FA patients, consistent with the critical role of retinoic acid metabolism and signaling during early development (14,21). FA patients frequently present with heterogenous multisystemic congenital defects, including short stature, microcephaly, microphthalmia, and upper and lower limb defects. The FA-D2 (*FANCD2^-/-^*) cells described in this study were isolated from a 1-year-old patient with a particularly severe clinical phenotype, characterized by multiple congenital defects (Supplementary Table 1), in addition to hypersensitivity to DEB-induced chromosome breakage during diagnosis (A. Smogorzewska, personal communication). Our findings suggest that dysregulated retinoic acid metabolism and signaling during development could play a significant role in the etiology of FA patient congenital abnormalities.

While important roles for the FA proteins in the repair of DNA interstrand crosslinks (ICLs) generated by mitomycin C and cisplatin have been clearly defined *in vitro*, the endogenous sources of genome instability in the physiological setting remain poorly understood. Over the past decade, several studies have established a critical role for certain FA proteins in a 2-tier system designed to mitigate the cytotoxic and genotoxic effects of reactive metabolic aldehydes, including acetaldehyde and formaldehyde (9,10). For example, *Aldh2^-/-^ Fancd2^-/-^* mice exhibit congenital defects and increased risk of leukemia, while aged *Aldh2^-/-^ Fancd2^-/-^* mice that escape leukemia develop aplastic anemia (5,6). Similarly, *Aldh2* gene inactivation or inhibition is synthetic lethal with loss of the homologous recombination and FA proteins Brca1, Brca2, and Rad51 (22). A synthetic lethal interaction between the alcohol dehydrogenase 5 gene *ADH5*, which restricts the cellular accumulation of formaldehyde, and the FA pathway has also been established (7). Our study indicates that the FA pathway also plays an important role in the mitigation of retinaldehyde genotoxicity. Cellular exposure to retinaldehyde resulted in the induction of phosphorylation of histone variant H2AX, a marker of DNA double-strand breaks (DSBs), albeit to a lesser extent than the topoisomerase type II inhibitor etoposide. In contrast to etoposide, which led to a marked increase in CHK1 S345 phosphorylation, indicative of activation of the intra-S-phase checkpoint, retinaldehyde exposure primarily induced phosphorylation of CHK2 T68, which is associated with activation of the G2-M checkpoint. We observed increased phosphorylation of H2AX and CHK2 T68 in both FA-A (*FANCA^-/-^*) and FA- D2 (*FANCD2^-/-^*) cells treated with retinaldehyde compared to FANCA- or FANCD2- complemented cells, respectively. Furthermore, following exposure to retinaldehyde, we observed increased *γ*H2AX nuclear foci formation in FA-D2 (*FANCD2^-/-^*) cells compared to FANCD2-complemented cells. Combined, these results indicate that the FA pathway plays an important role in the repair of retinaldehyde-induced DNA damage. Our results also suggest that certain cell lineages that are particularly dependent on retinoic acid signaling, e.g., neural stem and progenitor cells and hematopoietic stem cells, may be acutely reliant on an intact FA DNA repair pathway.

ALDH1A1 is one of 19 evolutionarily conserved human aldehyde dehydrogenase (ALDH) enzymes that catalyze the NAD(P)^+^-dependent irreversible oxidation of aldehydes to their corresponding carboxylic acids (23). The importance of the *ALDH* gene family is underscored by their association with increased risk for numerous pathologies including Sjögren-Larsson syndrome, type II hyperprolinemia, alcohol-related diseases, cancer, and neurodegeneration (23,24). The ALDH1A subfamily, comprised of ALDH1A1, ALDH1A2, and ALDH1A3, all exhibit specificity for retinaldehyde, with ALDH1A1 exhibiting a preference for 9-*cis*-retinaldehyde while ALDH1A3 exhibits a preference for all-*trans*-retinaldehyde (25,26). In our studies, we observed higher levels of phosphorylated H2AX following exposure to all-*trans*- retinaldehyde compared to 9-*cis*-retinaldehyde. Immunohistology studies have detected increased ALDH1A1 levels in a wide spectrum of cancers, with increased ALDH1A1 levels frequently associated with a poorer prognosis (24,27). The mechanisms by which ALDH1A1 promotes cancer cell stemness and poorer outcomes in cancer patients remain to be clearly elucidated. While we observed evidence for increased DNA damage and genotoxicity in retinaldehyde-treated FA cells, we did not observe differential sensitivity to retinaldehyde cytotoxicity between patient and complemented FA lines (results not shown). We speculate that increased levels of ALDH1A1 lead to increased oxidation of retinaldehyde to the biologically active molecule retinoic acid, which plays a key role in the transcriptional regulation of a wide variety of biological processes including proliferation, differentiation, and apoptosis (12,21). Indeed, all-*trans*-retinoic acid is used to rescue arrested differentiation in *PML-RARA* acute promyelocytic leukemia (APL) (28). Consistent with this prediction, we observed differential expression of numerous retinoic acid-regulated genes in non-transformed FA-D2 (*FANCD2^-/-^*) patient cells, e.g., *RARA*, *EGR1*, and *TGM2*, and several members of the *HOX*, *PAX*, and *SOX* gene families (results not shown). In our study, gene set enrichment analysis revealed differential expression of apoptosis genes between FA-D2 (*FANCD2^-/-^*) cells and the FANCD2- complemented cells. Attenuated apoptosis in FA-D2 (*FANCD2^-/-^*) cells was functionally validated using immunofluorescence microscopy for cleaved-PARP1 and immunoblotting for cleaved-PARP1 and cleaved caspase 3 and caspase 7. Taken together, our findings indicate that altered retinoic acid metabolism and signaling is likely to modulate several pathways and endpoints in non-transformed FA-D2 (*FANCD2^-/-^*) cells.

A previous study established an important role for the FA genes *BRCA1/FANCS* and *PALB2/FANCN* in the transcriptional regulation of retinoic acid-regulated genes (29). Depletion of *BRCA1/FANCS* and *PALB2/FANCN* resulted in decreased expression of the *HOXA1* and *HOXA2* genes following treatment with retinoic acid (29). BRCA1/FANCS and PALB2/FANCN were shown to be recruited to the promoters of retinoic acid-regulated genes indicating that these proteins may play a direct role in the transcriptional regulation of retinoic acid-responsive genes (29). Our studies establish an important link between retinoic acid metabolism - including retinaldehyde detoxification and retinoic acid-mediated transcriptional regulation - and the FA pathway. Considering the critical role of retinoic acid during embryogenesis, altered retinoic acid metabolism may contribute to many aspects of the pathophysiology of FA including the heterogeneous congenital abnormalities and increased risk for bone marrow failure observed among FA patients. Nervous system defects have become increasingly prevalent among FA patients (30,31,32). While retinoic acid is a well-established and critical signaling molecule during embryonal neuronal patterning and differentiation, retinoic acid also plays a key role in adult axon outgrowth and nerve regeneration and in the maintenance of the differentiated state of adult neurons (33,34). Further studies using developmentally appropriate and/or tissue-specific models are necessary to further define the role of retinoic acid metabolism in the pathophysiology of FA.

## MATERIALS AND METHODS

### Cell culture and generation of cell lines

The non-transformed FA-D2 (*FANCD2^-/-^*) patient samples described in this manuscript originated from a male patient diagnosed at the Rockefeller University International Fanconi Anemia Registry (IFAR) (Supplementary Table 1). Blood lymphocytes were collected at 1.5 months of age and EBV-immortalized (RA2472). Skin fibroblasts (RA2645) were prepared at 1 year of age. Fetal cells (RA2683) are from a female affected sibling miscarriage. These cells harbor a maternally inherited missense *FANCD2* hypomorphic mutation leading to an R815Q change and paternally inherited 2715+1G>A frameshift that leads to a truncated protein. Skin fibroblasts RA2645 were complemented with wild-type FANCD2 and functionally characterized by Helmut Hanenberg at the University of Duisberg-Essen and Detlev Schlinder at the University of Wuerzburg. FA-D2 (*FANCD2^-/-^*) and FANCD2-complemented cells were immortalized with human telomerase and grown in DMEM complete medium supplemented with 1 µg/ml puromycin. HSC72 (FA-A) and FANCA-complemented, EUFA121 (FA-F) and FANCF-complemented, and PD20 (FA-D2) and FANCD2-complemented patient-derived EBV-immortalized lymphoblasts were grown in Roswell Park Memorial Institute 1640 medium (RPMI 1640) supplemented with 20% (vol/vol) fetal bovine serum (FBS), 2 mM L-glutamine, and 50 µM penicillin-streptomycin. HSC72 (FA- A) lymphoblasts were derived from a 10-year-old female patient harboring a homozygous in-frame deletion of exons 18-28 in the *FANCA* gene. EUFA121 (FA-F) lymphoblasts were derived from a male patient of unknown age with compound heterozygous mutations in the *FANCF* gene (c.16C>T, c.351_397del) (35). PD20 cells were derived from a 7-year-old male patient with compound heterozygous mutations in the *FANCD2* gene (c.376A>G, c.3707G>A) (36). Mouse ear fibroblasts, A549 lung carcinoma, HeLa cervical carcinoma, Hepa1c1c7 mouse liver hepatoma, and U2OS osteosarcoma cells were grown in Dulbecco’s modified Eagle’s medium (DMEM) supplemented with 10% (vol/vol) FBS, L-glutamine, and penicillin-streptomycin.

### Immunoblotting

For immunoblotting analysis cell pellets were washed in phosphate-buffered saline (PBS) and lysed in 2% (wt/vol) SDS, 50 mM Tris-HCl, and 10 mM EDTA followed by sonication for 10s at 10% amplitude using a Sonic Dismembrator model 500 (Fisher Scientific). Proteins were resolved on NuPAGE 3-8% (wt/vol) Tris-acetate or 4-12% (wt/vol) Bis-Tris gels (Invitrogen) and transferred to polyvinylidene difluoride (PVDF) membranes. The following antibodies were used: rabbit polyclonal antisera against AKR1C3 (11194-1-AP; Proteintech), ALDH2 (18818S; Cell Signaling), ATR pS428 (2853S; Cell Signaling), CRABP2 (10225-1-AP; Proteintech), FANCD2 (NB100-182; Novus Biologicals), MRE11 (PC388; Calbiochem), PARP (9542; Cell Signaling), and RDH10 (14644-1-AP; Proteintech), or rabbit monoclonal antisera against ALDH1A1 (54135; Cell Signaling), ATM pS1981 (5883; Cell Signaling), ATR (13934; Cell Signaling), Caspase-3 (14220; Cell Signaling), Caspase-7 (12827; Cell Signaling), CHK1 pS345 (2348T; Cell Signaling), CHK2 pT68 (2197; Cell Signaling), Cleaved Caspase-3 (D175) (9664; Cell Signaling), Cleaved Caspase-7 (D198) (8438; Cell Signaling), Cleaved PARP (D214) (5625; Cell Signaling), and H2AX pS139 (*γ*H2AX) (9718; Cell Signaling), or mouse monoclonal antisera against alpha-tubulin (MS-581-PO; Neomarkers), ATM (11G12) (92356; Cell Signaling), CHK1 (2G1D5) (2360; Cell Signaling), p53 (DO-1) (F0909; Santa Cruz Biotechnology), p53 pS15 (9286; Cell Signaling), and RPA2 (NA18; Calbiochem).

### ALDEFLUOR assay

Cells were counted and adjusted to a concentration of 5 x 10^5^ cells/mL with ALDEFLUOR assay (01700; Stem Cell Technologies) buffer. Activated ALDEFLUOR reagent was added to cell suspensions and mixed well. Subsequently, half of the cell suspension was transferred to tubes containing the aldehyde dehydrogenase inhibitor ALDEFLUOR N,N-diethylaminobenzaldehyde (DEAB). Experimental and control samples were incubated at 37°C for 45 minutes. Cells were washed with ALDEFLUOR buffer once and resuspended in ALDEFLUOR buffer for FACS analysis. Samples were analyzed on a BD Biosciences-LSRII flow cytometer.

### AlDeSense assay

Cells were plated and incubated at 37°C/5% CO_2_ overnight. The following day the growth media was removed, and cells were incubated with Opti-MEM supplemented with either control AlDeSense AM or AlDeSense AM probes (20). Cells were incubated at room temperature with AlDeSense probes for 30 minutes and then imaged with a Zeiss Axio Observer A.1 inverted fluorescent microscope with ZEN lite image acquisition software.

### RNA sequencing

Quadruplicate biological replicates of non-transformed FA-D2 (*FANCD2^-/-^*) and FANCD2-complemented FA-D2 cells were harvested and washed with ice-cold PBS. Cell pellets were flash frozen and sent for sequencing with GENEWIZ/Azenta Life Sciences. RNA library prep was completed *via* polyA selection and HiSeq sequencing. RNA samples were quantified using Qubit 2.0 Fluorometer (Life Technologies, Carlsbad, CA) and RNA integrity was checked using Agilent TapeStation 4200 (Agilent Technologies, Palo Alto, CA). Sequencing libraries were prepared using NEBNext Ultra II RNA library prep kit using the manufacturer’s instructions (NEB, Ipswich, MA). Sequencing libraries were validated on the Agilent TapeStation and quantified using Qubit 2.0 Fluorometer as well as via quantitative PCR (KAPA Biosystems, Wilmington, MA). Sequencing libraries were clustered on a single lane of a flowcell and loaded onto an Illumina HiSeq 4000 instrument. Samples were sequenced using a 2 x150 bp paired-end configuration at 60 million reads per sample.

### RNA-seq analysis

Reads were trimmed to remove adapter sequences and bases of poor quality using Trimmomatic v.0.36. Trimmed reads were then mapped to the hg19 ENSEMBL reference genome using STAR aligner v.2.5.2b. FeatureCounts subread package v.1.5.2 was run on BAM files generated from the alignment, to calculate reads uniquely mapping to genes in the genome. FeatureCounts output files were used for downstream differential expression analysis using DESeq2.

### Immunofluorescence microscopy

For immunofluorescence microscopy (IF) analysis, cells were seeded in 8-well tissue culture slides (Millipore) and treated with retinaldehyde for 2 h, followed by a recovery period of 0, 3, or 6 h, or with staurosporine for 3 h only. For combined FANCD2 and *γ*H2AX IF, cells were fixed in 4% w/v paraformaldehyde and 2% w/v sucrose at 4°C followed by permeabilization in 0.3% v/v Triton X-100 in PBS. For cleaved PARP IF, cells were fixed in ice-cold 100% methanol. Fixed cells were blocked for 30 mins in antibody dilution buffer (5% v/v goat serum, 0.1% v/v NP40 in PBS) followed by incubation with anti-FANCD2 (NB100-182, Novus Biologicals) and anti-H2AX pS139 (JBW301, Millipore) or cleaved PARP (D214) (5625; Cell Signaling) antibodies for 1 h. Cells were washed three times in PBS and incubated in goat anti-mouse Texas Red (T-862, ThermoFisher Scientific) and goat anti-rabbit Fluorescein (F-2765, ThermoFisher Scientific) secondary antibody for 1 hour at room temperature. The slides were counterstained and mounted in vectashield plus 4’6-diamidine-2- phenylindole dihydrochloride (DAPI). Nuclear foci were analyzed using a Zeiss Axioimager.A2 upright epifluorescence microscope with ZEN lite image acquisition software.

### G2/M accumulation assay

Cells were seeded in 10 cm^2^ dishes and incubated in the absence or presence of mitomycin C (MMC) or etoposide (VP-16) for 24 h, or retinaldehyde (RALD) for 2 h, released and allowed to recover for 0, 3, 6, and 24 h. Cells were harvested, resuspended in PBS and fixed in ice-cold methanol. Samples were then washed in PBS and incubated with 50 µg/ml propidium iodide (PI) (Sigma) and 1x RNase A for 15 min at 37°C, followed by analysis using a BD FACSVerse flow cytometer. The percentages of cells in G_1_, S, and G_2_/M were determined by analyzing PI histograms with Flowjo v10.7.1 software.

### Crystal violet cell viability assay

Cells were seeded in 6-well dishes at a concentration of 40,000 cells/well and allowed to adhere at 37°C/5% CO_2_ overnight. Cells were then incubated with drug for 72 h at 37°C. After treatment, the media was aspirated, cells were washed once with PBS, and a 0.5% w/v crystal violet solution was added to each well. Plates were subsequently washed in water, allowed to dry overnight, and imaged. Finally, 100% methanol was added to each well and incubated for 2-3 h at 4°C. Methanol solution was then transferred to 96-well plates where the optical density of each well was measured at 570 nm (OD _570_). The average OD_570_ of non-treated cells was set to 100% and viability of treated cells were compared to cells that weren’t exposed to drug.

### siRNA

Cells were plated at 700,000 cells per 10 cm^2^ dish. The following day cells were transfected with SMARTpool siALDH1A1 (Horizon Discovery), siRDH10 (Horizon Discovery), or a control non-targeting siRNA using lipofectamine 2000 (ThermoFisher Scientific). Cells were incubated with siALDH1A1, siRDH10, or siControl for 24 h followed by a 24 h recovery period. Cells were harvested 96 h post treatment, washed with ice cold PBS, and used for downstream experiments.

### Gene set enrichment analysis

Gene sets enriched in either FA-D2 (*FANCD2^-/-^*) or FANCD2- complemented FA-D2 cells were analyzed by gene set enrichment analysis (GSEA) (v4.2.1.) software sourced by the Broad Institute (https://www.gsea-msigdb.org/gsea/index.jsp). Four gene sets including Angiogenesis (HALLMARK_ANGIOGENESIS.v7.5.1.gmt), Neurogenesis (GOBP_NEUROGENESIS.v7.5.1.gmt), Cellular Response to Retinoic Acid (GOBP_CELLULAR_RESPONSE_TO_RETINOIC_ACID.v7.5.1.gmt), and Apoptosis (HALLMARK_APOPTOSIS.v2022.1.Hs.gmt) were utilized on normalized read counts of each sequenced sample. Gene sets tested were considered to be significantly enriched in either FANCD2-complemented FA-D2 or FA-D2 (*FANCD2^-/-^*) phenotype at a nominal P value < 0.01.

### DNA extraction and mutation analysis

DNA was extracted from cells using DNA Blood and Tissue kit (Qiagen). PCR primers were designed using IDT Primer Quest to amplify regions around the mutation sites. Allele 1 Forward primer: 5’CAC GTT GTT GAG GAC AGT TCT A 3’, Reverse primer: 5’ CAG GGA TAT TGG CCT GAG ATT T 3’. Allele 2 Forward primer: 5’ CCT CTT CTA CCT CTA GGC AGT T 3’, Reverse primer: 5’ CCT TCA CGG ATA CCC ATC ATT C 3’. PCR was performed using Phusion HF Mastermix (ThermoFisher). Amplicons were cleaned with NucleoMag beads (New England Biolabs) and amplification was verified with gel electrophoresis. Sanger sequencing was performed on an ABI 3500xl instrument and raw data was analyzed using a mixed base detection protocol with a mixed base being called if the 2nd highest peak is at least 25% of the primary peak.

## ACKNOWLEDGEMENTS

We would like to express our sincere gratitude to the FA patients and their families for donating tissue to the International Fanconi Anemia Registry (IFAR) at the Rockefeller University. We thank A. Auerbach and A. Smogorzewska at the IFAR/Rockefeller University for providing deidentified patient material and clinical information. We thank H. Hanenberg at the University of Duisberg-Essen for complementation vectors. We thank the Fanconi Anemia Research Fund and Paul R. Andreassen at Cincinnati Children’s Hospital for providing cells. We thank J. Chan at the Beckman Institute for Advanced Science and Technology at the University of Illinois for providing the AlDeSense reagents. We thank members of the N. Howlett, G. Sun, J. Camberg, and C. Fallini laboratories for helpful discussions. This work was supported by an American Society of Hematology Bridge Grant to N.G.H., National Institutes of Health/National Heart, Lung, and Blood Institute R01 HL149907 to N.G.H., and Rhode Island IDeA Network of Biomedical Research Excellence (RI-INBRE) grant P20GM103430 from the National Institute of General Medical Sciences (PI, B.P. Cho).

## LEGENDS TO FIGURES

**Figure S1. Differential expression of Aldh1a1 and Rdh10 in non-transformed FA cell models.** (A) ALDH1A1, RDH10, and CRABP2 expression are largely confined to the cytosol. Non-transformed FA-D2 (*FANCD2^-/-^*) and FANCD2-complemented cells were fractionated into soluble and chromatin-associated fractions and lysates were immunoblotted with the indicated antibodies. Tubulin acts as a positive control for soluble proteins while histone H2A acts as a positive control for chromatin-associated proteins. (B) Immunoblotting analysis of EBV- immortalized FA-A (*FANCA^-/-^*) and FANCA-complemented, FA-F (*FANCF^-/-^*) and FANCF- complemented, and FA-D2 (*FANCD2^-/-^*) and FANCD2-complemented patient lymphoblasts using A549 and non-transformed FA-D2 (*FANCD2^-/-^*) and FANCD2-complemented fibroblasts as positive and negative controls. Whole-cell lysates were prepared and immunoblotted with the indicated antibodies. (B) Immunoblotting analysis of Fancd2^+/+^/Aldh2^+/+^, Fancd2^+/+^/Aldh2^-/-^, Fancd2^-/-^/Aldh2^+/+^, and Fancd2^-/-^/Aldh2^-/-^ mouse ear fibroblasts using A549, Hepa1c1c7, and non-transformed FA-D2 (*FANCD2^-/-^*) and FANCD2-complemented fibroblasts as positive and negative controls. *, Non-specific band; ALDH1A1*, ALDH1A1 immunoreactive band. (C) Immunoblotting analysis of Fancd2^+/+^, Fancd2^+/-^, and Fancd2^-/-^ mouse embryonic fibroblasts (MEFs) using A549, Hepa1c1c7, and non-transformed FA-D2 (*FANCD2^-/-^*) and FANCD2- complemented fibroblasts as positive and negative controls.

**Figure S2. Increased retinaldehyde genotoxicity in FA-D2 (FANCD2-/-) and FA-A (FANCA- /-) patient cells.** HeLa (A) and U2OS (B) cells were exposed to either 25 μM all-*trans*-retinaldehyde (a*t*RALD) or 25 μM 9-*cis-*retinaldehyde (9*c*RALD) for 2 h, released into drug-free media, and whole-cell lysates were prepared at 0, 3, 6, and 24 h following the exposure period. Whole-cell lysates were immunoblotted with the indicated antibodies. (C) FA-D2 (*FANCD2^-/-^*) and FANCD2-complemented cells were exposed to 25 μM a*t*RALD for 2 h and released into drug-free media. Cells were harvested at the indicated times, fixed, stained with propidium iodide, and analyzed by flow cytometry. (D) FA-D2 (*FANCD2^-/-^*) and FANCD2-complemented cells were exposed to 25 μM a*t*RALD or 25 μM VP-16 for 2 h and released into drug-free media. Immunofluorescence microscopy for *γ*H2AX was performed at 0, 6, and 24 h following release. Shown here are representative images of nuclei from FA-D2 (*FANCD2^-/-^*) and FANCD2- complemented cells 24 h following release from a*t*RALD and VP-16 treatment. (E) EBV- immortalized FA-A (*FANCA^-/-^*) and FANCA-complemented lymphoblasts were exposed to various concentrations of mitomycin C (MMC) for 48 h and percent survival determined using the MTT assay. Errors bars represent the standard deviations from 6 biological replicate measurements from a single experiment. This experiment was repeated at least three times with similar findings. (F) The same cells were exposed to 25 μM all-*trans-*retinaldehyde (a*t*RALD) for 2 h, released into a*t*RALD-free media, and whole-cell lysates were prepared at 0, 3, and 6 h following the exposure period. Whole-cell lysates were immunoblotted with the indicated antibodies.

## REFERENCES

1. Farf I. (2014) Fanconi Anemia: Guidelines for Diagnosis and Management. Fanconi Anemia Research Fund, Inc, Eugene, OR.

2. Auerbach A.D. and Wolman S.R. (1976) Susceptibility of Fanconi’s anaemia fibroblasts to chromosome damage by carcinogens. Nature, 261, 494–496.

3. Auerbach A.D. (1993) Fanconi anemia diagnosis and the diepoxybutane (DEB) test. Exp. Hematol., 21, 731–733.

4. Kottemann M.C. and Smogorzewska A. (2013) 3700363; Fanconi anaemia and the repair of Watson and Crick DNA crosslinks. Nature, 493, 356–363.

5. Langevin F., Crossan G.P., Rosado I.V., Arends M.J. and Patel K.J. (2011) Fancd2 counteracts the toxic effects of naturally produced aldehydes in mice. Nature, 475, 53–58.

6. Garaycoechea J.I., Crossan G.P., Langevin F., Daly M., Arends M.J. and Patel K.J. (2012) Genotoxic consequences of endogenous aldehydes on mouse haematopoietic stem cell function. Nature, 489, 571–575.

7. Rosado I.V., Langevin F., Crossan G.P., Takata M. and Patel K.J. (2011) Formaldehyde catabolism is essential in cells deficient for the Fanconi anemia DNA-repair pathway. Nat Struct Mol Biol, 18, 1432–1434.

8. Pontel L.B., Rosado I.V., Burgos-Barragan G., Garaycoechea J.I., Yu R., Arends M.J., Chandrasekaran G., Broecker V., Wei W., Liu L. et al. (2015) Endogenous formaldehyde is a hematopoietic stem cell genotoxin and metabolic carcinogen. Mol. Cell, 60, 177–188.

9. Langevin F.P., Garaycoechea J.I., Crossan G.P. and Patel K.J. (2013) [Aldehydes and Fanconi anaemia: The enemy within]. Med Sci (Paris), 29, 361–364.

10. Garaycoechea J.I. and Patel K.J. (2014) Why does the bone marrow fail in Fanconi anemia? Blood, 123, 26–34.

11. Vilhais-Neto G.C. and Pourquié O. (2008) Retinoic acid. Curr. Biol., 18, 191.

12. Blomhoff R. and Blomhoff H.K. (2006) Overview of retinoid metabolism and function. J. Neurobiol., 66, 606–630.

13. Bastos Maia S., Rolland Souza A.S., Costa Caminha M.F., Lins da Silva S., Callou Cruz, R. S. B. L., Carvalho Dos Santos C. and Batista Filho M. (2019) Vitamin A and pregnancy: A narrative review. Nutrients, 11, 681. doi: 10.3390/nu11030681.

14. Berenguer M. and Duester G. (2022) Retinoic acid, RARs and early development. J. Mol. Endocrinol., 69, T59–T67.

15. Walden H. and Deans A.J. (2014) The Fanconi anemia DNA repair pathway: Structural and functional insights into a complex disorder. Annu Rev Biophys, 43, 257–278.

16. Kalb R., Neveling K., Hoehn H., Schneider H., Linka Y., Batish S.D., Hunt C., Berwick M., Callen E., Surralles J. et al. (2007) 1852747; Hypomorphic mutations in the gene encoding a key fanconi anemia protein, FANCD2, sustain a significant group of FA-D2 patients with severe phenotype. Am. J. Hum. Genet., 80, 895-910.

17. Garcia-Higuera I., Taniguchi T., Ganesan S., Meyn M.S., Timmers C., Hejna J., Grompe M. and D’Andrea A.D. (2001) Interaction of the Fanconi anemia proteins and BRCA1 in a common pathway. Mol. Cell, 7, 249–262.

18. Thul P.J., Åkesson L., Wiking M., Mahdessian D., Geladaki A., Ait Blal H., Alm T., Asplund A., Björk L., Breckels L.M. et al. (2017) A subcellular map of the human proteome. Science, 356, eaal3321. doi: 10.1126/science.aal3321. Epub 2017 May 11.

19. Uhlén M., Fagerberg L., Hallström B.M., Lindskog C., Oksvold P., Mardinoglu A., Sivertsson Å, Kampf C., Sjöstedt E., Asplund A. et al. (2015) Proteomics. tissue-based map of the human proteome. Science, 347, 1260419.

20. Anorma C., Hedhli J., Bearrood T.E., Pino N.W., Gardner S.H., Inaba H., Zhang P., Li Y., Feng D., Dibrell S.E. et al. (2018) Surveillance of cancer stem cell plasticity using an isoform-selective fluorescent probe for aldehyde dehydrogenase 1A1. ACS Cent. Sci., 4, 1045–1055.

21. Das B.C., Thapa P., Karki R., Das S., Mahapatra S., Liu T.C., Torregroza I., Wallace D.P., Kambhampati S., Van Veldhuizen P. et al. (2014) Retinoic acid signaling pathways in development and diseases. Bioorg. Med. Chem., 22, 673–683.

22. Tacconi E.M., Lai X., Folio C., Porru M., Zonderland G., Badie S., Michl J., Sechi I., Rogier M., Matía García V. et al. (2017) BRCA1 and BRCA2 tumor suppressors protect against endogenous acetaldehyde toxicity. EMBO Mol. Med., 9, 1398–1414.

23. Jackson B., Brocker C., Thompson D.C., Black W., Vasiliou K., Nebert D.W. and Vasiliou V. (2011) Update on the aldehyde dehydrogenase gene (ALDH) superfamily. Hum. Genomics, 5, 283–303.

24. Marcato P., Dean C.A., Giacomantonio C.A. and Lee P.W.K. (2011) Aldehyde dehydrogenase: Its role as a cancer stem cell marker comes down to the specific isoform. Cell. Cycle, 10, 1378–1384.

25. Tomita H., Tanaka K., Tanaka T. and Hara A. (2016) Aldehyde dehydrogenase 1A1 in stem cells and cancer. Oncotarget, 7, 11018–11032.

26. Ma I. and Allan A.L. (2011) The role of human aldehyde dehydrogenase in normal and cancer stem cells. Stem Cell. Rev. Rep., 7, 292–306.

27. Yue H., Hu Z., Hu R., Guo Z., Zheng Y., Wang Y. and Zhou Y. (2022) ALDH1A1 in cancers: Bidirectional function, drug resistance, and regulatory mechanism. Front. Oncol., 12, 918778.

28. Warrell R.P.J., Frankel S.R., Miller W.H.J., Scheinberg D.A., Itri L.M., Hittelman W.N., Vyas R., Andreeff M., Tafuri A. and Jakubowski A. (1991) Differentiation therapy of acute promyelocytic leukemia with tretinoin (all-trans-retinoic acid). N. Engl. J. Med., 324, 1385–1393.

29. Gardini A., Baillat D., Cesaroni M. and Shiekhattar R. (2014) Genome-wide analysis reveals a role for BRCA1 and PALB2 in transcriptional co-activation. EMBO J., 33, 890–905.

30. Johnson-Tesch B.A., Gawande R.S., Zhang L., MacMillan M.L. and Nascene D.R. (2017) Fanconi anemia: Correlating central nervous system malformations and genetic complementation groups. Pediatr. Radiol., 47, 868–876.

31. Stivaros S.M., Alston R., Wright N.B., Chandler K., Bonney D., Wynn R.F., Will A.M., Punekar M., Loughran S., Kilday J.P. et al. (2015) Central nervous system abnormalities in Fanconi anaemia: Patterns and frequency on magnetic resonance imaging. Br. J. Radiol., 88, 20150088.

32. Aksu T., Gümrük F., Bayhan T., Coşkun Ç, Oğuz K.K. and Unal S. (2020) Central nervous system lesions in fanconi anemia: Experience from a research center for Fanconi anemia patients. Pediatr. Blood Cancer., 67, e28722.

33. Jacobs S., Lie D.C., DeCicco K.L., Shi Y., DeLuca L.M., Gage F.H. and Evans R.M. (2006) Retinoic acid is required early during adult neurogenesis in the dentate gyrus. Proc. Natl. Acad. Sci. U. S. A., 103, 3902–3907.

34. Maden M. (2007) Retinoic acid in the development, regeneration, and maintenance of the nervous system. Nat. Rev. Neurosci., 8, 755–765.

35. de Winter J.P., van Der Weel L., de Groot J., Stone S., Waisfisz Q., Arwert F., Scheper R.J., Kruyt F.A., Hoatlin M.E. and Joenje H. (2000) The Fanconi anemia protein FANCF forms a nuclear complex with FANCA, FANCC and FANCG. Hum.Mol.Genet., 9, 2665–2674.

36. Timmers C., Taniguchi T., Hejna J., Reifsteck C., Lucas L., Bruun D., Thayer M., Cox B., Olson S., D’Andrea A. et al. (2001) Positional cloning of a novel Fanconi anemia gene, FANCD2. Mol. Cell, 7, 241-248.

