## Supplementary figures and images for "Differential Regulation of Retinoic Acid Metabolism in Fanconi Anemia"

### Supplemental Figure 1

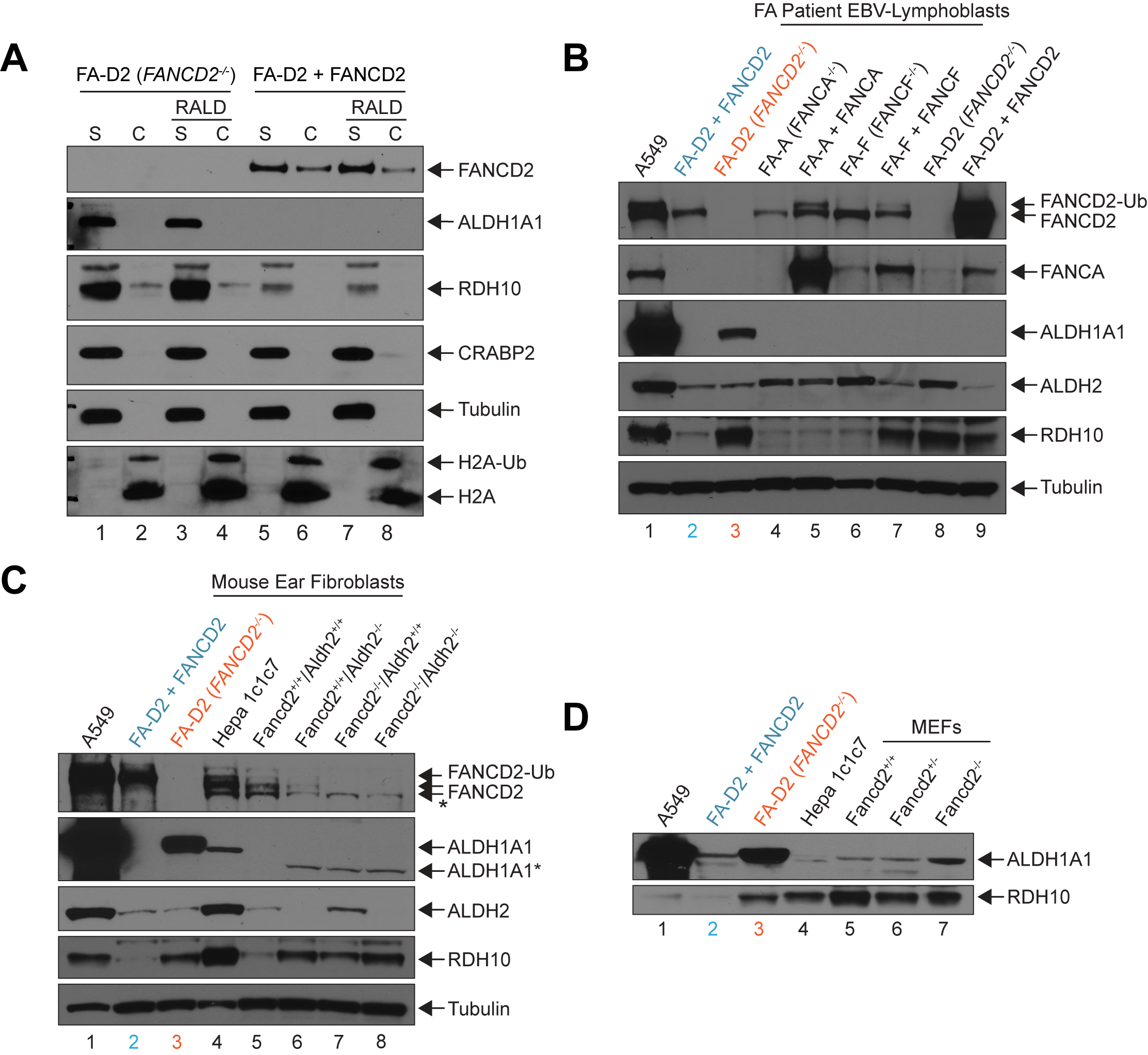

### Supplemental Figure 2A and B

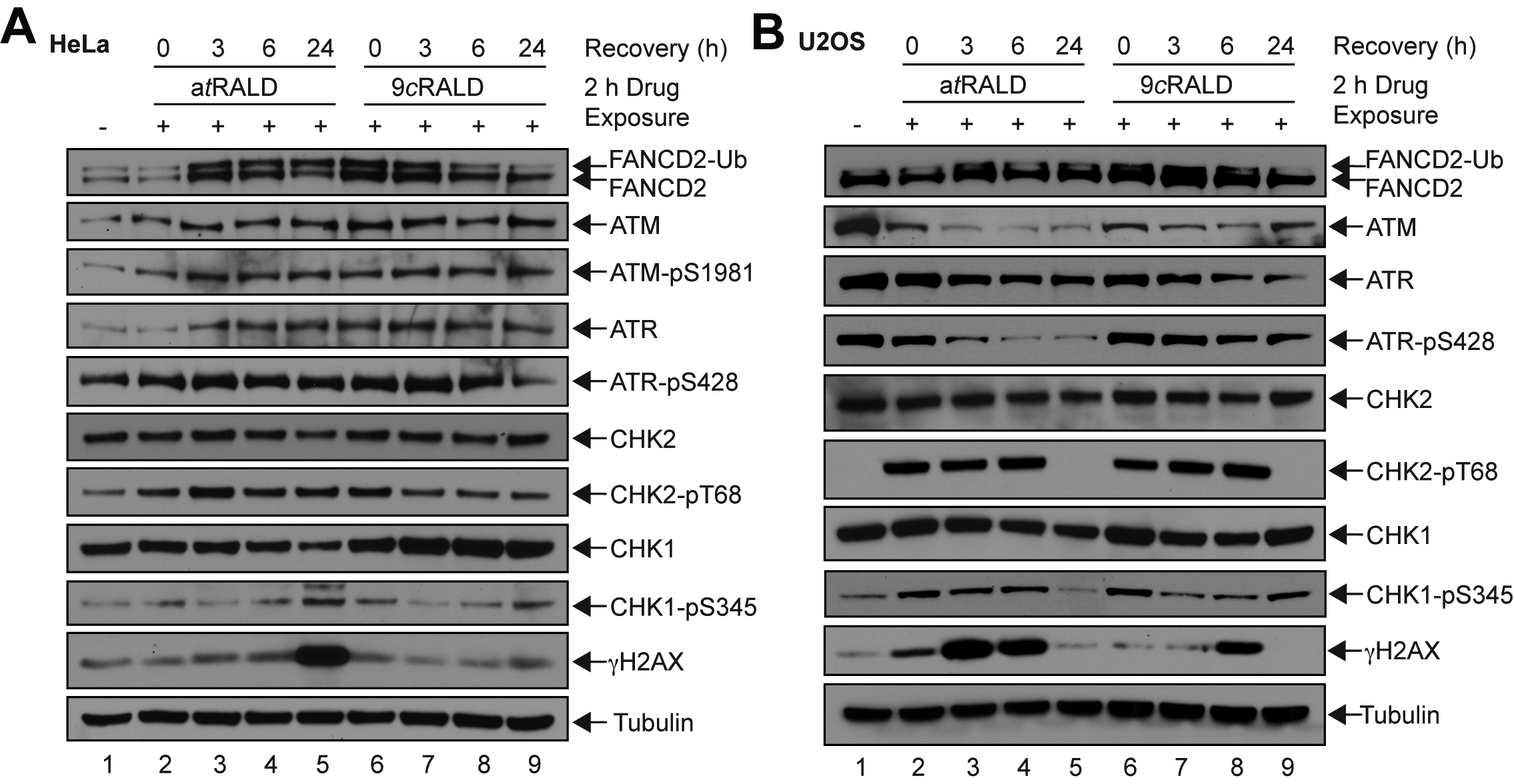

### Supplemental Figure 2C

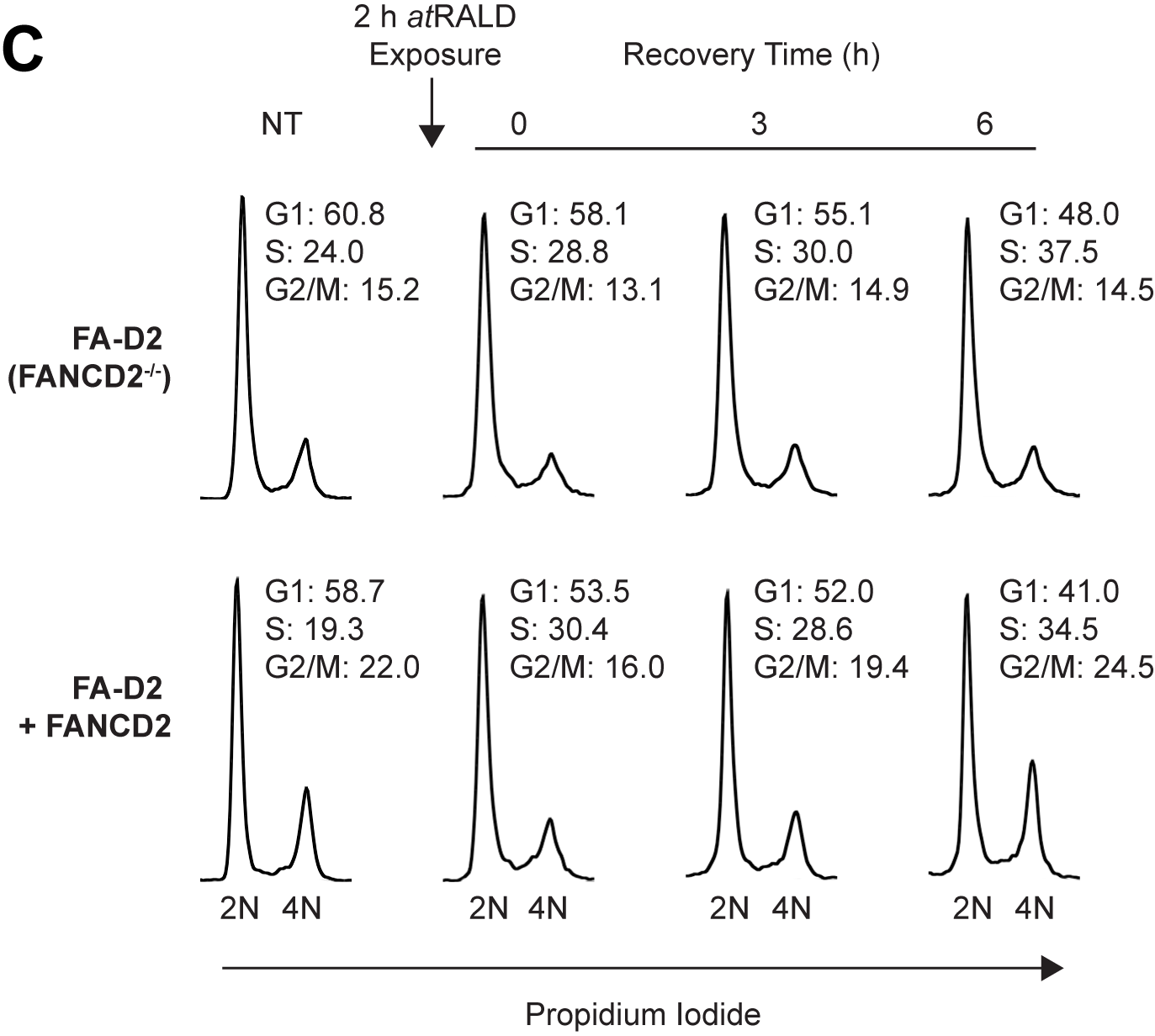

### Supplemental Figure 2D

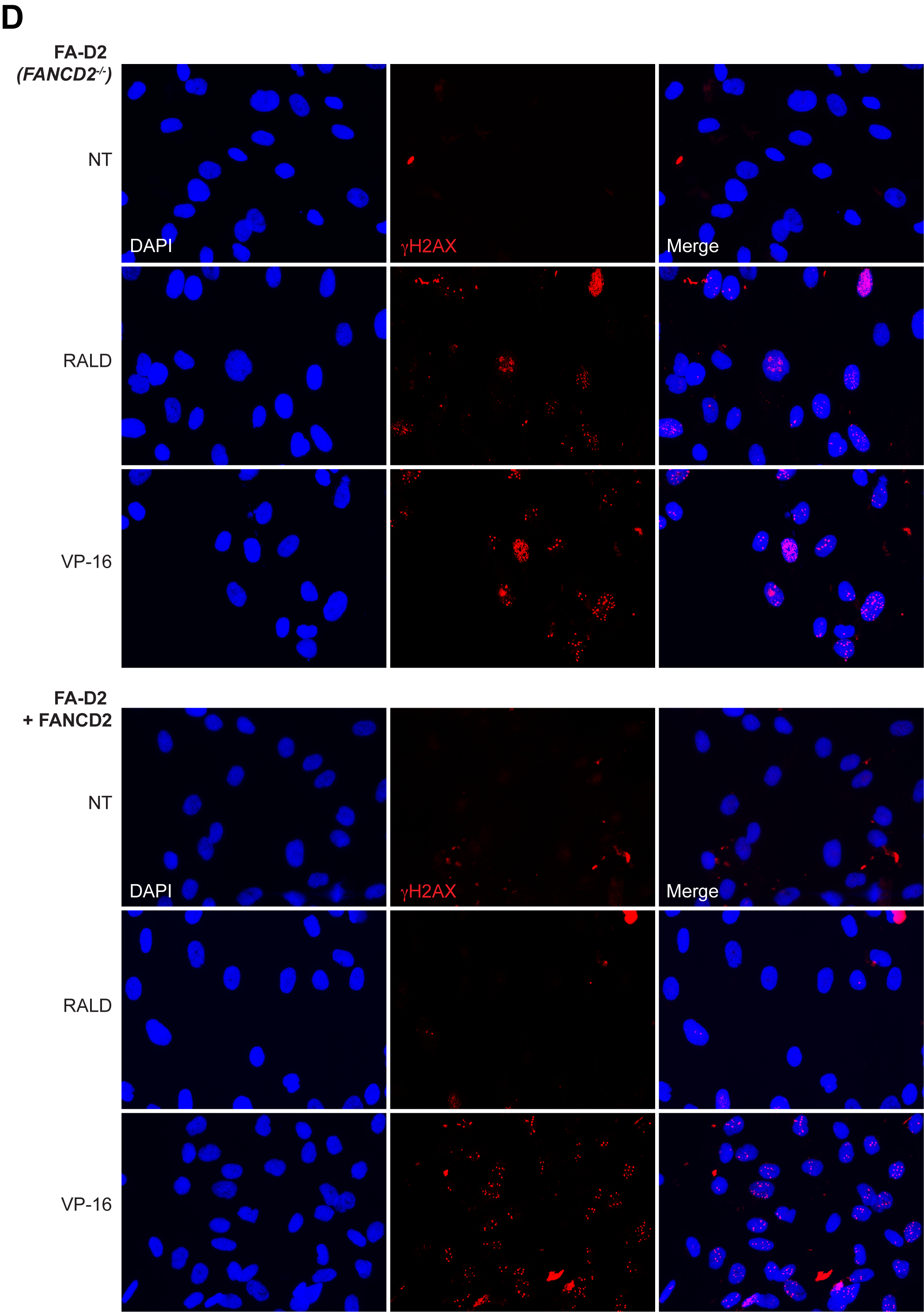

### Supplemental Figure 2E and F

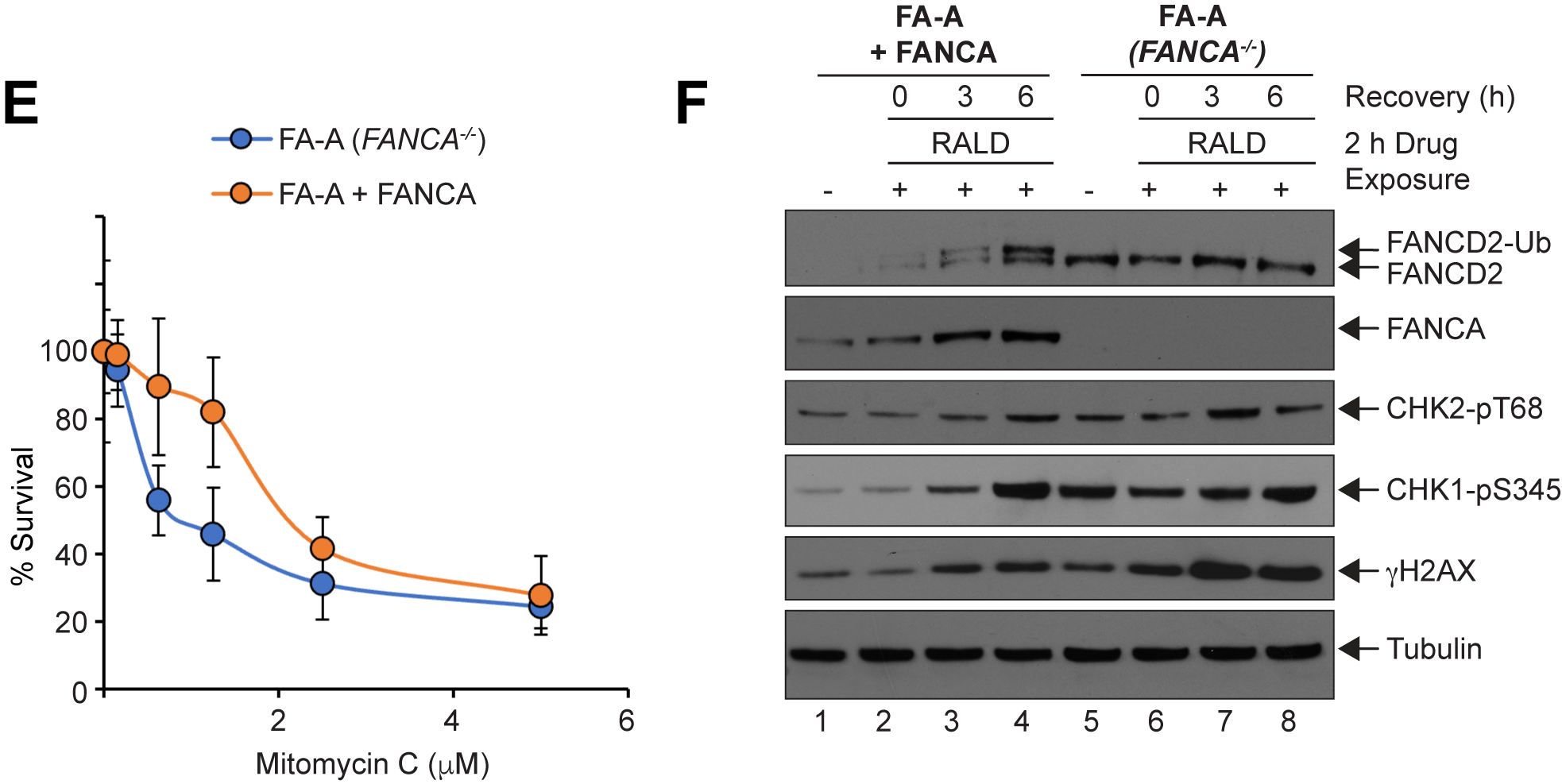
