## Supplemental Table 1 for "Differential Regulation of Retinoic Acid Metabolism in Fanconi Anemia"

**Supplementary Table 1.** Mutational and clinical characteristics of FA-D2 (*FANCD2^-/-^*) patient RA2427/RA2645.

| **Patient Sample ID** | **FA Complementation Group** | **Mutations** | **Clinical Presentation** |
| --- | --- | --- | --- |
| RA2427 (EBV-immortalized lymphoblasts)  RA2645 (Primary skin fibroblasts) | FA-D2 | *Allele 1*  c.2444G>A (exon 26)  r.2444G>A  p.(R815Q )  *Allele 2*  c.2715+1G>A  (IVS28+1G>A)  r.2715_2716ins27 (aberrant splicing),  p.(E906Lfs*4)** | - Mild frontal bossing - Café-au-lait spots - C4-C7 vertebral fusion - Hypoplastic thumbs (bilateral); Floating left thumb, Polydactyly of right thumb - Thenar hypoplasia (left) - High arched palate - Hypoplasia of the auditory canal (bilateral) - Low set ears (bilateral) - Microcephaly - Osteopenia - Constriction band on mid-forearms (bilateral) - Patent ductus arteriosus - Patent foramen ovale - Proportionate short stature - Retrognathia - Sacral dimple - Dystopia canthorum - Thin vermillion border - Velopharyngeal insufficiency |

**, While this allele is predicted to lead to the production of a truncated protein, a truncated FANCD2 isoform was not detected by western blotting.
