## Supplemental Table 2 for "Differential Regulation of Retinoic Acid Metabolism in Fanconi Anemia"

**Supplementary Table 2.** Average RNA-seq normalized read counts of retinoic acid metabolism and signaling genes differentially expressed in FA-D2 (*FANCD2^-/-^*) patient cells and FA-D2 (*FANCD2^-/-^*) cells complemented with wild-type FANCD2.

| **Gene name** | **Ensembl ID** | **Log2FC** | ***P*adj** | **FA-D2 (*FANCD2^-/-^*)** | **FA-D2 + FANCD2** |
| --- | --- | --- | --- | --- | --- |
| *ALDH1A1* | ENSG00000165092 | -7.2 | 0.000 | 1134.3 ± 147.4 | 7.5 ± 6.6 |
| *ADH1B* | ENSG00000196616 | -2.4 | 0.001 | 2662.1 ± 767.9 | 516.2 ± 357.5 |
| *SLC6A4* | ENSG00000108576 | -1.7 | 0.019 | 16.8 ± 7.9 | 5.3 ± 1.7 |
| *RDH10* | ENSG00000121039 | -1.4 | 0.000 | 1250.7 ± 383.0 | 482.8 ± 34.5 |
| *LTK* | ENSG00000062524 | -1.4 | 0.015 | 19.4 ± 3.6 | 7.4 ± 1.8 |
| *WNT11* | ENSG00000085741 | -1.2 | 0.020 | 34.6 ± 7.1 | 15.0 ± 7.3 |
| *CRABP2* | ENSG00000143320 | -1.1 | 0.000 | 19840.7 ± 3322.6 | 9550.4 ± 979.9 |
| *CYP26B1* | ENSG00000003137 | -1.1 | 0.020 | 792.9 ± 419.7 | 381.7 ± 119.7 |
| *AQP1* | ENSG00000240583 | 1.0 | 0.000 | 1205.7 ± 196.9 | 2429.2 ± 273.5 |
| *RARA* | ENSG00000131759 | 1.0 | 0.000 | 1304.1 ± 359.8 | 2662.8 ± 261.9 |
| *LYN* | ENSG00000254087 | 1.1 | 0.001 | 36.9 ± 10.0 | 77.1 ± 4.8 |
| *PTCH1* | ENSG00000185920 | 1.1 | 0.009 | 105.6 ± 56.6 | 225.9 ± 45.8 |
| *MYB* | ENSG00000118513 | 1.3 | 0.025 | 8.4 ± 4.0 | 21.4 ± 4.6 |
| *TNC* | ENSG00000041982 | 1.9 | 0.000 | 11678.1 ± 1950.5 | 42967.8 ± 4369.0 |
| *COL1A1* | ENSG00000108821 | 2.0 | 0.000 | 302817.7 ± 71696.4 | 1189028.0 ± 451874.5 |
| *AKR1C3* | ENSG00000196139 | 2.3 | 0.000 | 134.5 ± 54.2 | 655.9 ± 178.9 |
| *LEP* | ENSG00000174697 | 2.5 | 0.031 | 3.6 ± 4.0 | 20.6 ± 23.2 |
| *BRINP1* | ENSG00000078725 | 2.5 | 0.000 | 14.0 ± 2.6 | 80.0 ± 11.9 |
| *CYP1A1* | ENSG00000140465 | 2.7 | 0.000 | 15.0 ± 8.5 | 101.3 ± 23.5 |
