## Supplemental Table 3 for "Differential Regulation of Retinoic Acid Metabolism in Fanconi Anemia"

**Supplementary Table 3.** Average RNA-seq normalized read counts of the 19 human aldehyde dehydrogenase genes in FA-D2 (*FANCD2^-/-^*) patient cells and FA-D2 (*FANCD2^-/-^*) cells complemented with wild-type FANCD2.

| **Gene name** | **Ensembl ID** | **Log2FC** | ***P*adj** | **FA-D2 (*FANCD2^-/-^*)** | **FA-D2 + FANCD2** |
| --- | --- | --- | --- | --- | --- |
| *ALDH1A1* | ENSG00000165092 | -7.2 | 8.2 x 10^-92^ | 1134.3 ± 147.4 | 7.5 ± 6.6 |
| *ALDH1A2* | ENSG00000128918 | ne | ne | ne | ne |
| *ALDH1A3* | ENSG00000184254 | -0.7 | 0.015 | 5809.9 ± 1779.4 | 3481.0 ± 761.7 |
| *ALDH1B1* | ENSG00000137124 | -0.1 | 0.603 | 2001.6 ± 314.1 | 1848.0 ± 200.2 |
| *ALDH1L1* | ENSG00000144908 | ne | ne | ne | ne |
| *ALDH1L2* | ENSG00000136010 | 0.9 | 0.001 | 2583.8 ± 393.7 | 4731.4 ± 1358.3 |
| *ALDH2* | ENSG00000111275 | -0.6 | 0.000 | 1451.5 ± 182.5 | 961.1 ± 66.9 |
| *ALDH3A1* | ENSG00000108602 | 1.0 | 0.003 | 134.8 ± 41.1 | 273.3 ± 86.1 |
| *ALDH3A2* | ENSG00000072210 | -0.6 | 0.004 | 4487.4 ± 296.6 | 2925.4 ± 640.8 |
| *ALDH3B1* | ENSG00000006534 | 0.2 | 0.103 | 2917.5 ± 190.5 | 3451.8 ± 322.9 |
| *ALDH3B2* | ENSG00000132746 | ne | ne | ne | ne |
| *ALDH4A1* | ENSG00000159423 | -0.2 | 0.426 | 1446.2 ± 255.4 | 1281.5 ±126.6 |
| *ALDH5A1* | ENSG00000112294 | -0.5 | 0.480 | 19.6 ± 11.8 | 13.9 ± 3.0 |
| *ALDH6A1* | ENSG00000119711 | -0.6 | 0.003 | 1320.7 ± 191.9 | 869.5 ± 124.4 |
| *ALDH7A1* | ENSG00000164904 | -0.4 | 0.046 | 3909.2 ± 651.0 | 3048.8 ± 64.7 |
| *ALDH8A1* | ENSG00000118514 | ne | ne | ne | ne |
| *ALDH9A1* | ENSG00000143149 | -0.5 | 0.014 | 4995.2 ± 1098.6 | 3508.7 ± 71.4 |
| *ALDH16A1* | ENSG00000161618 | 0.1 | 0.734 | 1658.5 ± 384.7 | 1779.8 ± 325.4 |
| *ALDH18A1* | ENSG00000059573 | -0.1 | 0.851 | 10471.8 ± 2328.8 | 10103.3 ± 986.8 |
