## Supplemental Table 4 for "Differential Regulation of Retinoic Acid Metabolism in Fanconi Anemia"

**Supplementary Table 4.** Average RNA-seq normalized read counts of human alcohol dehydrogenase *ADH* genes in FA-D2 (*FANCD2^-/-^*) patient cells and FA-D2 (*FANCD2^-/-^*) cells complemented with wild-type FANCD2.

| **Gene name** | **Ensembl ID** | **Log2FC** | ***P*adj** | **FA-D2 (*FANCD2^-/-^*)** | **FA-D2 + FANCD2** |
| --- | --- | --- | --- | --- | --- |
| *ADH1B* | ENSG00000196616 | -2.4 | 0.001 | 2662.1 ± 767.9 | 516.2 ± 357.5 |
| *ADH1C* | ENSG00000248144 | -1.3 | 0.050 | 20.8 ± 5.5 | 8.1 ± 4.9 |
| *ADH5* | ENSG00000197894 | -1.1 | 0.000 | 17197.4 ± 3789.1 | 8298.1 ± 702.7 |
| *ADHFE1* | ENSG00000147575 | -0.5 | 0.148 | 65.0 ± 16.2 | 46.1 ± 4.5 |
